# Plankton food webs of the Gulf of Mexico spawning grounds of Atlantic Bluefin tuna

**DOI:** 10.1101/2020.07.29.227116

**Authors:** Michael R. Stukel, Trika Gerard, Thomas Kelly, Angela N. Knapp, Raúl Laiz-Carrión, John Lamkin, Michael R. Landry, Estrella Malca, Karen E. Selph, Akihiro Shiroza, Taylor A. Shropshire, Rasmus Swalethorp

## Abstract

We used linear inverse ecosystem modeling techniques to assimilate data from extensive Lagrangian field experiments into a mass-balance constrained food web for the Gulf of Mexico open-ocean ecosystem. This region is highly oligotrophic, yet Atlantic Bluefin Tuna (ABT) travel long distances from feeding grounds in the North Atlantic to spawn there. Our results show that the food web is dominated by the microbial loop (>80% of net primary productivity is respired by heterotrophic bacteria and protists that feed on them). In contrast, herbivorous food web pathways from phytoplankton to metazoan zooplankton process <4% of net primary production in the mixed layer. Nevertheless, ABT larvae feed preferentially on calanoid copepods and other suspension-feeding zooplankton that in turn derive much of their nutrition from diatoms and mixotrophic flagellates. This allows ABT larvae to maintain a comparatively low trophic level (∼4.0 for pre-flexion larvae; ∼4.2 for post-flexion larvae) that increases trophic transfer from phytoplankton to larval fish.

## INTRODUCTION

The open-ocean Gulf of Mexico (GoM) is a nutrient-poor, low plankton biomass region (Biggs & Ressler, 2001, Damien *et al.*, 2018, Muller-Karger *et al.*, 2015, Shropshire *et al.*, 2020). Nevertheless, it is an important region for spawning and larval development of many commercially-important fishes (Cornic *et al.*, 2018, Kitchens & Rooker, 2014, Lindo-Atichati *et al.*, 2012, Rooker *et al.*, 2013, Rooker *et al.*, 2012). The western stock of Atlantic Bluefin Tuna (ABT) travel long distances from feeding grounds throughout the North Atlantic to spawning grounds in the oligotrophic GoM, implying that some characteristics of this region enhance larval success (Rooker *et al.*, 2007, Teo *et al.*, 2007). One strong possibility is that low predator abundances in this food-poor region are a prerequisite for pelagic larvae to survive to maturity (Biggs, 1992). Nevertheless, it remains unclear how ABT and other GoM larval fishes manage to obtain sufficient nutrition during their critical first-feeding period.

Discerning the structure of GoM planktonic food webs is crucial to answering such questions. ABT larvae are selective feeders that rely disproportionately on specific prey taxa including calanoid and poecilostomatoid copepods, cladocerans, and appendicularians (Llopiz *et al.*, 2015, Llopiz *et al.*, 2010, Shiroza *et al.*, in prep., Tilley *et al.*, 2016). These prey items, however, have distinctly different trophic and ecological roles. Appendicularians are filter-feeding pelagic tunicates with fine meshes that give them access to some of the smallest cyanobacteria in the ocean (Alldredge, 1976, Gorsky & Fenaux, 1998). Poecilostomatoid copepods, by contrast, are predators of other metazoan zooplankton, and hence likely feed comparatively high on the food chain (Turner, 1986). Cladocerans and calanoid copepods are often omnivorous filter feeders, although calanoid copepods can fill multiple trophic roles within the planktonic food web (Bode *et al.*, 2015, Katechakis & Stibor, 2004, Mauchline *et al.*, 1998, Uye & Kayano, 1994).

Elucidating the linkages between larval fish, their prey, and the base of the planktonic food web is crucial to predicting climate change impacts on larval survival (Landry *et al.*, 2019). Different phytoplankton functional groups (e.g., *Prochlorococcus, Trichodesmium*, nanoflagellates, unicellular microbial diazotrophs) will have different responses to warming, acidification, and increased stratification in the oligotrophic ocean (Barton *et al.*, 2016, Flombaum *et al.*, 2013, Hong *et al.*, 2017, Rost *et al.*, 2008). These variable responses originate from different physiological responses to stressors, but also to fundamentally different relationships between these groups and limiting nutrient, light, or temperature conditions. For instance: *Trichodesmium* and other diazotrophs are not nitrogen limited, *Prochlorococcus* is well adapted to utilizing recycled nitrogen available at low concentrations in oligotrophic regions, and nanoflagellates may need to rely partially on phagotrophic behavior (mixotrophy) to alleviate nutrient stress (Scanlan & Post, 2008, Stoecker *et al.*, 2017, Zehr, 2011). The pathways that connect different nutrient sources (upwelling, lateral advection, recycled production, and diazotrophy) through phytoplankton and zooplankton to larval fishes will fundamentally determine how these organisms respond to climate change.

In this study, we use linear inverse models (LIM) as a data synthesis tool to constrain pelagic food webs of the oligotrophic GoM. We utilize results from field experiments designed to investigate the open-ocean GoM ecosystem from nutrients to fish (Landry & Lamkin, in prep.). LIM allows us to incorporate diverse ecosystem measurements (e.g., primary productivity, protistan grazing rates, copepod δ^15^N, and larval ABT gut contents) into a mass-balance constrained ecosystem model. We use the results to address three distinct questions: What is the trophic level of larval ABT and what is the trophic efficiency of food chains leading to larval ABT? Which phytoplankton groups ultimately support secondary production by larval ABT? Are specific food web pathways to larval ABT less reliant on upwelled nitrate?

## METHODS

### In situ measurements

Our data are derived from two cruises in ABT spawning grounds in April-May 2017 and 2018 as part of the Bluefin Larvae in Oligotrophic Ocean Foodwebs: Investigating Nutrients to Zooplankton in the Gulf of Mexico (BLOOFINZ-GoM) Project (Table 1). During these cruises, we conducted regional zooplankton sampling surveys, guided in part by the Bluefin Tuna index (Domingues et al., 2016), to identify contrasting open-ocean water parcels with and without high abundances of ABT larvae (Landry & Lamkin, in prep.). We then conducted three-to five-day Lagrangian experiments (hereafter “cycles”), while following satellite-enabled drift arrays with 3×1-m holey-sock drogues centered at 15 m depth (Landry, 2009, Stukel *et al.*, 2015).

**Table 1.** Rate, biomass, and δ^15^N measurements used as inputs to the inverse model.

| <u>Rate Measurements</u> | <u>Units</u> | <u>Cycle 1</u> | <u>Cycle 2</u> | <u>Cycle 3</u> | <u>Cycle 4</u> | <u>Cycle 5</u> | <u>Source</u> |
| --- | --- | --- | --- | --- | --- | --- | --- |
| NPP (shallow) | (mmol N m<sup>-2</sup> d<sup>-1</sup>) | 2.23 ± 0.13 | 1.93 ± 0.15 | 2.46 ± 0.26 | 2.4 ± 0.11 | 3.08 ± 0.14 | Yingling et al. (this iss |
| NPP (deep) | (mmol N m<sup>-2</sup> d<sup>-1</sup>) | 1.64 ± 0.07 | 1.75 ± 0.08 | 1.3 ± 0.08 | 1.92 ± 0.05 | 1.34 ± 0.05 | Yingling et al. (this iss |
| f-ratio (shallow) | (mmol N m<sup>-2</sup> d<sup>-1</sup>) | 0.06 ± 0.04 | 0.01 ± 0 | 0.04 ± 0.04 | 0.06 ± 0.02 | 0.14 ± 0.04 | Yingling et al. (this iss |
| f-ratio (deep) | (mmol N m<sup>-2</sup> d<sup>-1</sup>) | 0.44 ± 0.3 | 0.03 ± 0.01 | 0.07 ± 0.07 | 0.07 ± 0.02 | 0.09 ± 0.02 | Yingling et al. (this iss |
| Protistan Grazing Rate (shallow) | (mmol N m<sup>-2</sup> d<sup>-1</sup>) | 3.17 ± 1.11 | 2.22 ± 1.09 | 2.11 ± 0.57 | 2.76 ± 0.58 | 3.51 ± 0.19 | Yingling et al. (this iss |
| Protistan Grazing Rate (deep) | (mmol N m<sup>-2</sup> d<sup>-1</sup>) | 3.42 ± 1.15 | 2.68 ± 1.32 | 1.64 ± 0.47 | 2.78 ± 0.5 | 1.77 ± 0.49 | Yingling et al. (this iss |
| Picophyto NPP (shallow) | (mmol N m<sup>-2</sup> d<sup>-1</sup>) | 1.47 ± 0.23 | 0 ± 0 | 0 ± 0 | 0 ± 0 | 0 ± 0 | Landry et al. (this iss |
| Picophyto NPP (deep) | (mmol N m<sup>-2</sup> d<sup>-1</sup>) | 1.26 ± 0.22 | 0 ± 0 | 0 ± 0 | 0 ± 0 | 0 ± 0 | Landry et al. (this iss |
| Flagellate NPP (shallow) | (mmol N m<sup>-2</sup> d<sup>-1</sup>) | 2.79 ± 0.64 | 2.23 ± 1.75 | - | 1.41 ± 0.37 | 0.44 ± 0.17 | Landry et al. (this iss |
| Flagellate NPP (deep) | (mmol N m<sup>-2</sup> d<sup>-1</sup>) | 1.52 ± 0.76 | 0.7 ± 0.55 | - | 0.48 ± 0.2 | 0.77 ± 0.53 | Landry et al. (this iss |
| Diatom NPP (shallow) | (mmol N m<sup>-2</sup> d<sup>-1</sup>) | 0.16 ± 0.09 | 0.0028 ± 0.0034 | - | 0.03 ± 0.01 | 0.05 ± 0.03 | Landry et al. (this iss |
| Diatom NPP (deep) | (mmol N m<sup>-2</sup> d<sup>-1</sup>) | 0.15 ± 0.13 | 0.002 ± 0.0024 | - | 0.02 ± 0.01 | 0.03 ± 0.02 | Landry et al. (this iss |
| Picophyto Mortality (shallow) | (mmol N m<sup>-2</sup> d<sup>-1</sup>) | 1.01 ± 0.38 | 0 ± 0 | 0 ± 0 | 0 ± 0 | 0 ± 0 | Landry et al. (this iss |
| Picophyto Mortality (deep) | (mmol N m<sup>-2</sup> d<sup>-1</sup>) | 0.69 ± 0.06 | 0 ± 0 | 0 ± 0 | 0 ± 0 | 0 ± 0 | Landry et al. (this iss |
| Flagellate Mortality (shallow) | (mmol N m<sup>-2</sup> d<sup>-1</sup>) | 2.04 ± 0.35 | 1.73 ± 1.59 | - | 0.62 ± 0.14 | 0.6 ± 0.26 | Landry et al. (this iss |
| Flagellate Mortality (deep) | (mmol N m<sup>-2</sup> d<sup>-1</sup>) | 1.34 ± 0.41 | 0.19 ± 0.17 | - | 0.17 ± 0.17 | 0.87 ± 0.6 | Landry et al. (this iss |
| Diatom Mortality (shallow) | (mmol N m<sup>-2</sup> d<sup>-1</sup>) | 0.11 ± 0.06 | 0.001 ± 0.002 | - | 0.02 ± 0.01 | 0.04 ± 0.02 | Landry et al. (this iss |
| Diatom Mortality (deep) | (mmol N m<sup>-2</sup> d<sup>-1</sup>) | 0.02 ± 0.02 | 0.001 ± 0.001 | - | 0.01 ± 0 | 0.01 ± 0.01 | Landry et al. (this iss |
| NVM Mesozoo Grazing | (mmol N m<sup>-2</sup> d<sup>-1</sup>) | 0.39 ± 0.1 | 0.15 ± 0.02 | 0.46 ± 0.05 | 0.21 ± 0.04 | 0.52 ± 0.1 | Landry et al. (this iss |
| VM Mesozoo Grazing | (mmol N m<sup>-2</sup> d<sup>-1</sup>) | 0.03 ± 0.06 | 0.13 ± 0.06 | 0.09 ± 0.05 | 0.02 ± 0.02 | 0.15 ± 0.08 | Landry et al. (this iss |
| SedTrap Flux (shallow) | (mmol N m<sup>-2</sup> d<sup>-1</sup>) | 1.53 ± 0.55 | 0.82 ± 0.05 | 0.98 ± 0.26 | 0.59 ± 0.04 | 1.08 ± 0.07 | Kelly et al. (this iss |
| SedTrap Flux (deep) | (mmol N m<sup>-2</sup> d<sup>-1</sup>) | 0.46 ± 0.02 | 0.52 ± 0.18 | 1.14 ± 0.18 | 0.47 ± 0.1 | 0.87 ± 0.18 | Kelly et al. (this iss |
| Thorium Flux (shallow) | (mmol N m<sup>-2</sup> d<sup>-1</sup>) | - | - | - | - | - | Kelly et al. (this iss |
| Thorium Flux (deep) | (mmol N m<sup>-2</sup> d<sup>-1</sup>) | - | - | - | - | - | Kelly et al. (this iss |
| Chl Sinking (shallow) | (mmol N m<sup>-2</sup> d<sup>-1</sup>) | 0.02 ± 0.02 | 0.02 ± 0.01 | - | 0.01 ± 0 | 0.03 ± 0.02 | Kelly et al. (this iss |
| Chl Sinking (deep) | (mmol N m<sup>-2</sup> d<sup>-1</sup>) | 0.02 ± 0.01 | 0.06 ± 0.02 | - | 0.02 ± 0.02 | 0.05 ± 0.04 | Kelly et al. (this iss |
| Fecal Pellet Sinking (shallow) | (mmol N m<sup>-2</sup> d<sup>-1</sup>) | 0.03 ± 0.03 | 0.01 ± 0.01 | - | 0 ± 0 | 0.02 ± 0.02 | Kelly et al. (this iss |
| Fecal Pellet Sinking (deep) | (mmol N m<sup>-2</sup> d<sup>-1</sup>) | 0.13 ± 0.06 | 0.09 ± 0.06 | - | 0.14 ± 0.14 | 0.25 ± 0.2 | Kelly et al. (this iss |
| Microzoo to Preflex | (mmol N m<sup>-2</sup> d<sup>-1</sup>) | 0.16 ± 0.08 | - | - | - | - | Shiroza et al. (this iss |
| Microzoo to Postflex | (mmol N m<sup>-2</sup> d<sup>-1</sup>) | 2.06 ± 0.33 | - | - | - | - | Shiroza et al. (this iss |
| Appendicularian to Preflex | (mmol N m<sup>-2</sup> d<sup>-1</sup>) | 0.45 ± 0.18 | - | - | - | - | Shiroza et al. (this iss |
| Appendicularian to Postflex | (mmol N m<sup>-2</sup> d<sup>-1</sup>) | 3.27 ± 0.23 | - | - | - | - | Shiroza et al. (this iss |
| Cladoceran to Preflex | (mmol N m<sup>-2</sup> d<sup>-1</sup>) | 0.28 ± 0.28 | - | - | - | - | Shiroza et al. (this iss |
| Cladoceran to Postflex | (mmol N m<sup>-2</sup> d<sup>-1</sup>) | 22.03 ± 2.12 | - | - | - | - | Shiroza et al. (this iss |
| Calanoids to Preflex | (mmol N m<sup>-2</sup> d<sup>-1</sup>) | 5.46 ± 2.27 | - | - | - | - | Shiroza et al. (this iss |
| Calanoids to Postflex | (mmol N m<sup>-2</sup> d<sup>-1</sup>) | 37.72 ± 0.57 | - | - | - | - | Shiroza et al. (this iss |
| Poecilastomatoids to Preflex | (mmol N m<sup>-2</sup> d<sup>-1</sup>) | 0 ± 0.01 | - | - | - | - | Shiroza et al. (this iss |
| Poecilastomatoids to Postflex | (mmol N m<sup>-2</sup> d<sup>-1</sup>) | 8.47 ± 0.63 | - | - | - | - | Shiroza et al. (this iss |
| <u>**Biomass and other measurements**</u> |  |  |  |  |  |  |  |
| Temperature (0-50) | deg C | 24.31 | 25.16 | 26.18 | 25.22 | 24.44 | CTD |
| Temperature (50-120) | deg C | 22.14 | 23.87 | 23.96 | 24.4 | 21.68 | CTD |
| Temperature (120-300) | deg C | 16.41 | 19.28 | 18.99 | 18.91 | 16.46 | CTD |
| HerbNVM biomass (shallow) | ($\mu\text{mol N m}^{-2}$) | 8.94 | - | - | - | 45.44 | Shiroza et al. (this issue) |
| App Biomass (shallow) | ($\mu\text{mol N m}^{-2}$) | 0.22 | - | - | - | 0.32 | Shiroza et al. (this issue) |
| Clad Biomass (shallow) | ($\mu\text{mol N m}^{-2}$) | 14.91 | - | - | - | 18.13 | Shiroza et al. (this issue) |
| NVM Cal Biomass (shallow) | ($\mu\text{mol N m}^{-2}$) | 2.74 | - | - | - | 1.92 | Shiroza et al. (this issue) |
| Chaeto Biomass (shallow) | ($\mu\text{mol N m}^{-2}$) | 107.04 | - | - | - | 130.13 | Shiroza et al. (this issue) |
| Poecil Biomass (shallow) | ($\mu\text{mol N m}^{-2}$) | 10.82 | - | - | - | 11.07 | Shiroza et al. (this issue) |
| Gel Pred Biomass (shallow) | ($\mu\text{mol N m}^{-2}$) | - | - | - | - | - | |
| Preflex Biomass (shallow) | ($\mu\text{mol N m}^{-2}$) | 0.1 | - | - | - | - | Shiroza et al. (this issue) |
| Postflex Biomass (shallow) | ($\mu\text{mol N m}^{-2}$) | 1.12 | - | - | - | - | Shiroza et al. (this issue) |
| HerbVM Biomass | ($\mu\text{mol N m}^{-2}$) | 34.3 | - | - | - | - | Shiroza et al. (this issue) |
| vmCal Biomass | ($\mu\text{mol N m}^{-2}$) | 26.4 | - | - | - | - | Shiroza et al. (this issue) |
| HerbNVM biomass (deep) | ($\text{mmol N m}^{-2}$) | - | - | - | - | - | |
| App Biomass (deep) | ($\text{mmol N m}^{-2}$) | - | - | - | - | - | |
| Clad Biomass (deep) | ($\text{mmol N m}^{-2}$) | - | - | - | - | - | |
| NVM Cal Biomass (deep) | ($\text{mmol N m}^{-2}$) | - | - | - | - | - | |
| Chaeto Biomass (deep) | ($\text{mmol N m}^{-2}$) | - | - | - | - | - | |
| Poecil Biomass (deep) | ($\text{mmol N m}^{-2}$) | - | - | - | - | - | |
| Gel Pred Biomass (deep) | ($\text{mmol N m}^{-2}$) | - | - | - | - | - | |
| Cyano Biomass (shallow) | ($\text{mmol N m}^{-2}$) | 9.39 | 7.45 | 7.71 | 12.86 | 18.39 | Selph et al. (this issue) |
| Tricho Biomass (shallow) | ($\mu\text{mol N m}^{-2}$) | 27.23 | 5.15 | 17.05 | 3.5 | 0.97 | Selph et al. (this issue) |
| Diatom Biomass (shallow) | ($\text{mmol N m}^{-2}$) | 0.13 | 0.003 | 1.12 | 0.03 | 0.08 | Selph et al. (this issue) |
| Flag Biomass (shallow) | ($\text{mmol N m}^{-2}$) | 4.84 | 2.82 | 6.74 | 1.61 | 2.74 | Selph et al. (this issue) |
| HNF Biomass (shallow) | ($\text{mmol N m}^{-2}$) | - | - | - | - | - | |
| MIC Biomass (shallow) | ($\text{mmol N m}^{-2}$) | - | - | - | - | - | |
| Cyano Biomass (deep) | ($\text{mmol N m}^{-2}$) | 8.22 | 8.93 | 9 | 15.41 | 6.77 | Selph et al. (this issue) |
| Tricho Biomass (deep) | ($\mu\text{mol N m}^{-2}$) | 0.77 | 0.3 | 1.62 | 0.08 | 0.11 | Selph et al. (this issue) |
| Diatom Biomass (deep) | ($\text{mmol N m}^{-2}$) | 0.12 | -0.003 | -1.12 | 0.03 | 0.04 | Selph et al. (this issue) |
| Flag Biomass (deep) | ($\text{mmol N m}^{-2}$) | 6.97 | 8.97 | 5.43 | 1.63 | 3.63 | Selph et al. (this issue) |
| HNF Biomass (deep) | ($\text{mmol N m}^{-2}$) | - | - | - | - | - | |
| MIC Biomass (deep) | ($\text{mmol N m}^{-2}$) | - | - | - | - | - | |
| HerbNVM Size | ($\mu\text{g C ind}^{-1}$) | 1.35 | 1.35 | 1.35 | 1.35 | 1.35 | Shiroza et al. (this issue) |
| App Size | ($\mu\text{g C ind}^{-1}$) | 0.07 | 0.07 | 0.07 | 0.07 | 0.07 | Shiroza et al. (this issue) |
| Clad Size | ($\mu\text{g C ind}^{-1}$) | 2.63 | 2.63 | 2.63 | 2.63 | 2.63 | Shiroza et al. (this issue) |
| NVM Cal Size | ($\mu\text{g C ind}^{-1}$) | 4.44 | 4.44 | 4.44 | 4.44 | 4.44 | Shiroza et al. (this issue) |
| Chaeto Size | ($\mu\text{g C ind}^{-1}$) | 20.83 | 20.83 | 20.83 | 20.83 | 20.83 | Shiroza et al. (this issue) |
| Poecil Size | ($\mu\text{g C ind}^{-1}$) | 2.33 | 2.33 | 2.33 | 2.33 | 2.33 | Shiroza et al. (this issue) |
| Gel Pred Size | ($\mu\text{g C ind}^{-1}$) | - | - | - | - | - | |
| Preflex Size | ($\mu\text{g C ind}^{-1}$) | 83.71 | 83.71 | 83.71 | 83.71 | 83.71 | Laiz-Carrion et al. (this issue) |
| Postflex Size | ($\mu\text{g C ind}^{-1}$) | 179.75 | 179.75 | 179.75 | 179.75 | 179.75 | Laiz-Carrion et al. (this issue) |
| HerbVM Size | ($\mu\text{g C ind}^{-1}$) | 4.44 | 4.44 | 4.44 | 4.44 | 4.44 | Laiz-Carrion et al. (this issue) |
| vmCal Size | ($\mu\text{g C ind}^{-1}$) | 4.44 | 4.44 | 4.44 | 4.44 | 4.44 | Laiz-Carrión et al. (this issue) |
| Maximum upwelling rate (shallow) | ($\mu\text{mol N m}^{-2} \text{d}^{-1}$) | 0.09 | 0.09 | 0.09 | 0.09 | 0.09 | Kelly et al. (this issue) |
| Maximum upwelling rate (deep) | ($\mu\text{mol N m}^{-2} \text{d}^{-1}$) | 366.79 | 1142.9 | 1200.44 | 1020.28 | 1543.33 | Kelly et al. (this issue) |
| <u>$\delta^{15}\text{N}$ Values</u> |  |  |  |  |  |  |  |
| Upwelled Nitrate | $\delta^{15}\text{N}_{\text{AIR}} (\text{‰})$ | 3.20 | 3.10 | 2.80 | 2.00 | 2.90 | Knapp et al. (this issue) |
| Preflex ABT | $\delta^{15}\text{N}_{\text{AIR}} (\text{‰})$ | 4.63 | - | - | - | 7.50 | Swalethorp et al. (this issue) |
| Postflex ABT | $\delta^{15}\text{N}_{\text{AIR}} (\text{‰})$ | 4.21 | - | - | - | 6.16 | Swalethorp et al. (this issue) |
| Shallow SedTrap | $\delta^{15}\text{N}_{\text{AIR}} (\text{‰})$ | 2.90 | 2.51 | 1.68 | 2.50 | 3.80 | Kelly et al. (this issue) |
| Deep SedTrap | $\delta^{15}\text{N}_{\text{AIR}} (\text{‰})$ | 4.89 | 2.86 | 3.71 | 3.73 | 4.55 | Kelly et al. (this issue) |
| Shallow DON | $\delta^{15}\text{N}_{\text{AIR}} (\text{‰})$ | 3.37 | 3.37 | 3.37 | 3.45 | 3.27 | Knapp et al. (this issue) |
| Deep DON | $\delta^{15}\text{N}_{\text{AIR}} (\text{‰})$ | 3.31 | 3.31 | 3.31 | 3.33 | 3.39 | Knapp et al. (this issue) |
| Shallow PON | $\delta^{15}\text{N}_{\text{AIR}} (\text{‰})$ | 1.44 | 1.20 | 1.11 | 2.13 | 2.66 | Kelly et al. (this issue) |
| Deep PON | $\delta^{15}\text{N}_{\text{AIR}} (\text{‰})$ | 1.80 | 1.57 | 2.45 | 1.78 | 1.63 | Kelly et al. (this issue) |

During each cycle, we made daily profiles with a CTD-Niskin rosette to measure temperature, salinity, and density and collect samples for: chlorophyll *a* measurements (by the acidification method, Strickland & Parsons, 1972), phytoplankton pigment analyses (by high-pressure liquid chromatography), picophytoplankton and heterotrophic bacteria enumeration (by flow cytometry; Selph *et al.*, 2016, Selph *et al.*, in prep.), nano- and microphytoplankton biomass (Selph *et al.*, in prep., Taylor & Landry, 2018), *Trichodemium* biomass (Selph *et al.*, in prep.), nutrients (nitrate and ammonium; Knapp *et al.*, in prep.), dissolved organic nitrogen (DON, Knapp *et al.*, in prep.), particulate organic nitrogen (PON, Kelly *et al.*, in prep., Stukel *et al.*, 2016), and δ^15^N of nitrate, DON, and PON (Kelly *et al.*, in prep., Knapp *et al.*, 2005, Knapp *et al.*, in prep., Sigman *et al.*, 2001).

We also conducted a suite of daily in situ rate measurements that were incubated in mesh bags affixed at 6 depths (spanning the euphotic zone) on one of the floating arrays (Landry, 2009, Landry & Lamkin, in prep.). These measurements included nitrate uptake (Stukel *et al*., 2016, Yingling *et al.*, in prep.), net primary production (Yingling *et al*., in prep.), and group-specific phytoplankton growth and mortality due to protistan grazing (Landry *et al*., 2016, Landry *et al*., in prep.). All *in situ* incubations were conducted for 24 h at natural light and temperature conditions. We also conducted shorter (4 – 6 h) deckboard incubations for nitrate and ammonium uptake (Stukel *et al.*, 2016, Yingling *et al.*, in prep.).

Twice per day (mid-day and midnight) we conducted oblique net tows through the euphotic zone to collect mesozooplankton that were analyzed for dry weight, carbon, nitrogen, isotopes and gut pigment content (Décima *et al.*, 2019, Swalethorp *et al.*, in prep.). Gut pigment contents were analyzed as in Decima *et al.* (2016) to estimate grazing rates (Landry *et al.*, in prep.). More frequent net tows were conducted to a depth of 25 m to collect larval ABT for abundance, size, dry weight, gut content, otolith-based age, and isotopic measurements (Laiz-Carrion *et al.*, 2015, Malca *et al.*, 2017, Malca *et al.*, in prep., Shiroza *et al.*, in prep.).

Nitrogen inputs to and outputs from the euphotic zone were constrained using sediment traps, thorium measurements, and Thorpe scale analyses. Surface-tethered drifting sediment traps were used to collect sinking PON, chlorophyll, and phaeopigments at 50 m depth, near the base of the euphotic zone (∼120 m), and beneath the euphotic zone (200 m) (Kelly *et al.*, in prep., Stukel *et al.*, 2019). ^238^U-^234^Th disequilibrium was used as an independent estimate of sinking PON (Kelly *et al.*, in prep., Stukel *et al.*, 2019). We used Thorpe-scale analyses to constrain vertical eddy diffusivity and upward nitrate flux (Gargett & Garner, 2008, Kelly *et al.*, in prep.). We combined day-night differences in mesozooplankton biomass with allometric ammonium excretion relationships to quantify active transport by diel vertical migrants (Ikeda, 1985, Kelly *et al.*, in prep.).

### Food web structure

Our food web structure was specifically designed to address the variability in GoM food chains that channel energy towards the prey of ABT larvae (either efficiently or inefficiently) or to the multiple plankton taxa that are not suitable prey for ABT larvae (Fig. 1). The model includes three nutrient classes (NO_3_^-^, NH_4_^+^, and N_2_) and three non-living organic matter pools (DOM, small detritus, and large detritus). It includes four phytoplankton: *Trichodesmium* and unicellular cyanobacteria (which are assumed to be potentially diazotrophic), diatoms, and mixotrophic flagellates. It also includes heterotrophic bacteria, heterotrophic nanoflagellates, and microzooplankton. Six suspension-feeding mesozooplankton are included: appendicularians (which are the only suspension feeders capable of feeding on cyanobacteria and heterotrophic bacteria), vertically-migrating calanoid copepods, non-vertically migrating calanoid copepods, cladocerans, other non-vertically migrating herbivorous suspension feeders, and other vertically-migrating herbivorous suspension feeders. It includes two small predatory mesozooplankton: chaetognaths and poecilostomatoid copepods. It also includes 4 “higher trophic levels” that serve as closure terms in the model: preflexion ABT larvae, postflexion ABT larvae, other planktivorous fish, and predatory gelatinous zooplankton (e.g., ctenophores and cnidarians). Trophic pathways are determined based on known predator-prey relationships. Because ABT larvae feed only in the mixed layer, we include two layers in the model: mixed layer, and deep euphotic zone. All model compartments are identical, except that ABT larvae only exist in the mixed layer. The two layers are connected through upward flux of nitrate, downward flux of sinking particles and the motions of vertical-migratory taxa, which are assumed to be able to freely migrate between the two layers during the night, but reside beneath the euphotic zone (i.e., outside the model) during the day. Inputs to the model include upwelled nitrate, diazotrophy, and lateral advection of POM and DOM. Closure terms include secondary production of higher trophic levels, sinking of large detritus, sinking of diatoms, and sinking of mixotrophic flagellates, and excretion of vertical migratory taxa beneath the euphotic zone. We assume Redfield stoichiometry for all model flows, which allows us to relate respiration to ammonium excretion. We thus use the term “respiration” when relating respiratory or excretory fluxes to primary production and use the term “excretion” when discussing nutrient recycling. Supp. Table. 1 shows all model flows.

**Fig. 1.**
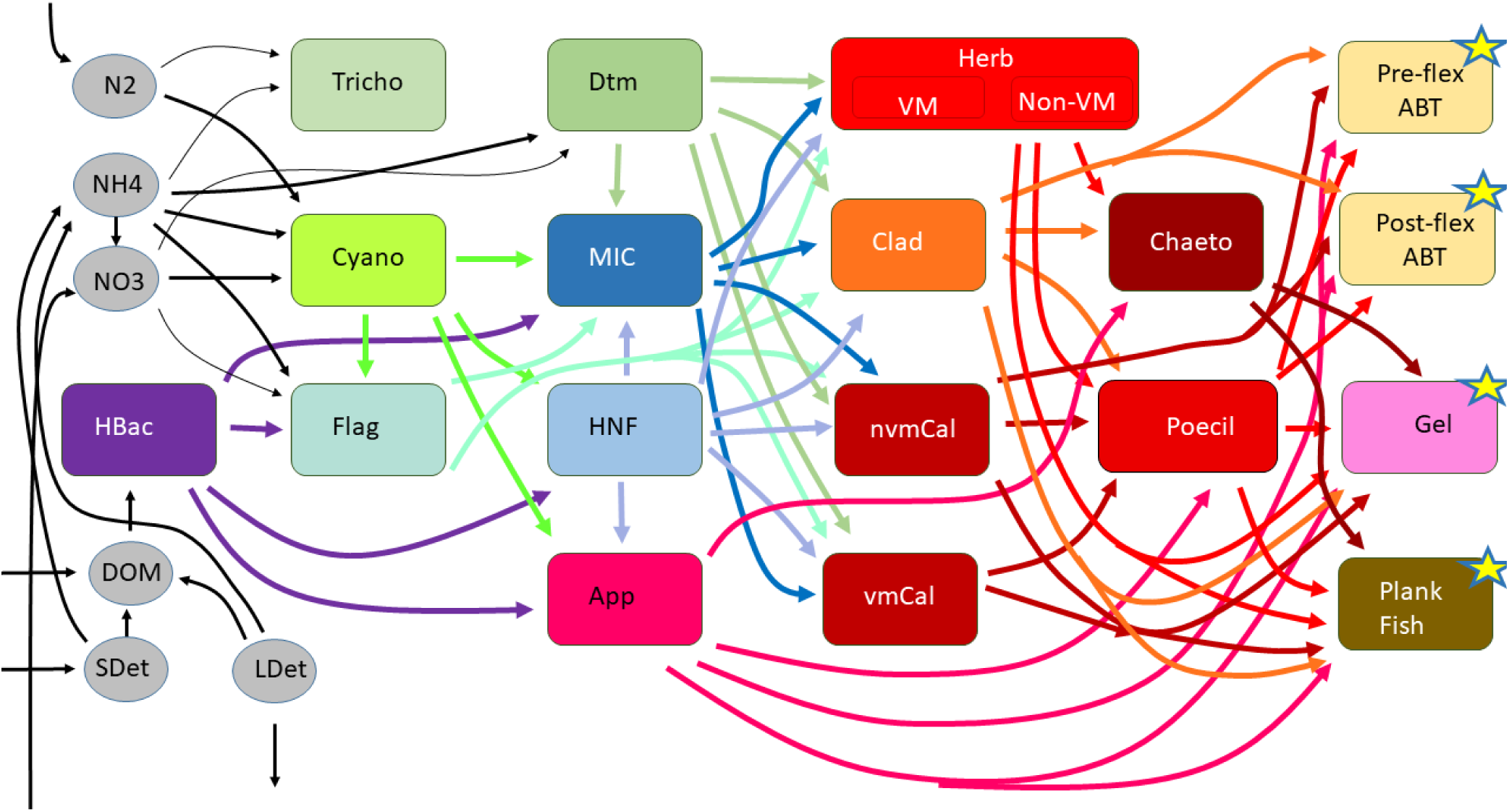
Food web structure. All major food web flows are shown (i.e., all flows between living organism groups). However, for visual simplicity, we omit production of NH_4_^+^, DOM, and detritus by all living groups as well as consumption of detritus by protistan zooplankton and suspension-feeding metazoans. Stars indicate the highest trophic levels in the model. The secondary production of these groups is a closure term in the model. The model utilizes a two-layer structure (∼mixed layer and deep euphotic zone). All trophic food web components exist in both layers except for larval ABT. For all model flows, see Supp. Table 1.

### Inverse model solution

To constrain the flux of nitrogen through unmeasured ecosystem pathways, we used linear inverse modeling (LIM) techniques (Van Oevelen *et al.*, 2010, Vézina & Platt, 1988). LIM allows investigators to specify mass balance constraints that must be exactly fit by food web solutions 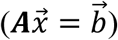, approximate equations that quantify measured rates with associated measurement uncertainty 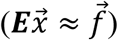, and inequality constraints 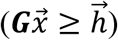 that represent *a priori* acceptable ranges for different ecosystem properties (e.g., gross growth efficiency of zooplankton is between 10% and 40%). We used a total of 44 mass-balance constraints (exact equalities), 82 *in situ* measurements (approximate equalities), and 248 inequality constraints. However, with 302 total unknown food web flows 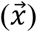, the system remains under-constrained. To objectively determine a representative solution (and confidence limits) we used the Markov Chain Monte Carlo (MCMC) with ^15^N approach (Stukel *et al.*, 2018a, Stukel *et al.*, 2018b). The MCMC approach initially uses the exact mass balance constraints to remove degrees of freedom from the solution and then creates bounds on the solution as formed by the hyperplanes prescribed by the inequality constraints (Kones *et al.*, 2009, Soetaert *et al.*, 2009, Van Den Meersche *et al.*, 2009). Then, starting with an initial guess of the solution that satisfies the equality and inequality constraints, the MCMC approach then conducts a random walk through the solution space bounded by 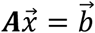 and 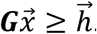. New solutions are accepted based on the relative misfits of the new and previous solution with respect to the approximate equality measurements 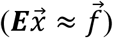. This generates a set of solutions satisfying the equality and inequality constraints, and with the probability of inclusion of a specific solution related to how well it satisfies the combined field measurement constraints. The mean solution of the MCMC approach has been shown to more accurately recover withheld measurement constraints than the previously used L_2_ minimum norm approach (Saint-Béat *et al.*, 2013, Stukel *et al.*, 2012). The MCMC+^15^N approach builds on this previous work, but allows for the incorporation of non-linear constraints associated with unknown δ^15^N values for some organisms or non-living nitrogen pools in the ecosystem. It uses a second varying solution vector 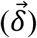 quantifying the ^15^N isotope fraction for each unknown nitrogen pool. A new solution set for 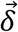 is determined at the same time as the new solution set for 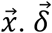 modifies approximate equality constraints associated with ^15^N mass balance. For additional details, see Stukel et al. (2018a).

### Food web analyses

Trophic levels (TL) for all zooplankton were computed as one plus the ingestion-weighted mean TL of prey. All phytoplankton were assumed TL=1, except mixotrophic flagellates, which had *TL* = (1 − *p*_*phag*_) + *p*_*phag*_(1 + *TL*_*prey*_), where p_phag_ is the proportion of their nitrogen derived from phagotrophy (rather than dissolved nutrient uptake). Heterotrophic bacteria were assumed to have a TL equal to one plus the TL of the organism producing the organic matter they utilized.

To quantify indirect nitrogen flows through the food web, we used indirect food web flow analysis (Hannon, 1973). The normalized amount of nitrogen (direct and indirect) that any organism derives from any other organism (or non-living nitrogen pool) can be computed as (*I* − *G*)^−1^, where I is the identity matrix and G is the normalized production matrix (i.e., a matrix giving the percentage of an organism’s nitrogen requirement derived from any other organism).

Following Stukel et al. (2012) we defined three major food web pathways that describe energy and nutrient fluxes from the base of the food web: the herbivorous food chain, the multivorous food chain, and the microbial loop. The herbivorous food chain was defined as the sum of nitrogen flux from phytoplankton to metazoan zooplankton. The multivorous food chain was defined as the sum of nitrogen flux that reaches metazoan zooplankton after passing through protistan grazers. The microbial loop was defined as the sum of bacterial respiration and the fraction of protistan respiration that was supported by bacterial production.

## RESULTS

### Model performance

The LIEM model did a decent job of matching field measurements. The square root mean squared error, which can be thought of as the average number of standard errors that model estimates were from the measurements, was 1.33 if we consider only the field rate measurements. It was 0.81 for all approximate equality equations (including the δ^15^N mass balance equations). The largest mismatches between model and measurements were with respect to growth and grazing estimates for specific phytoplankton taxa (particularly mixotrophic flagellates). These model-data misfits are driven in large part by inconsistency in the field measurements; the sum of phytoplankton production for all taxa as determined by the microzooplankton dilution approach was higher than primary production estimated using H^13^CO_3_^-^ uptake (Fig. 2). In comparison to rate measurements associated with lower trophic levels, the model very accurately recovered ingestion rates of larval ABT on different mesozooplankton groups (Fig. 2b). Model solutions were also strongly constrained by the comparatively low δ^15^N of larval ABT (Fig. 3). The model struggled to determine solution vectors that matched the comparatively low δ^15^N of larval ABT with the fairly similar measured δ^15^N of upwelled nitrate, sinking detritus, and bulk suspended organic matter, thus leading to model misfits in the δ^15^N mass balance equations.

**Fig. 2.**
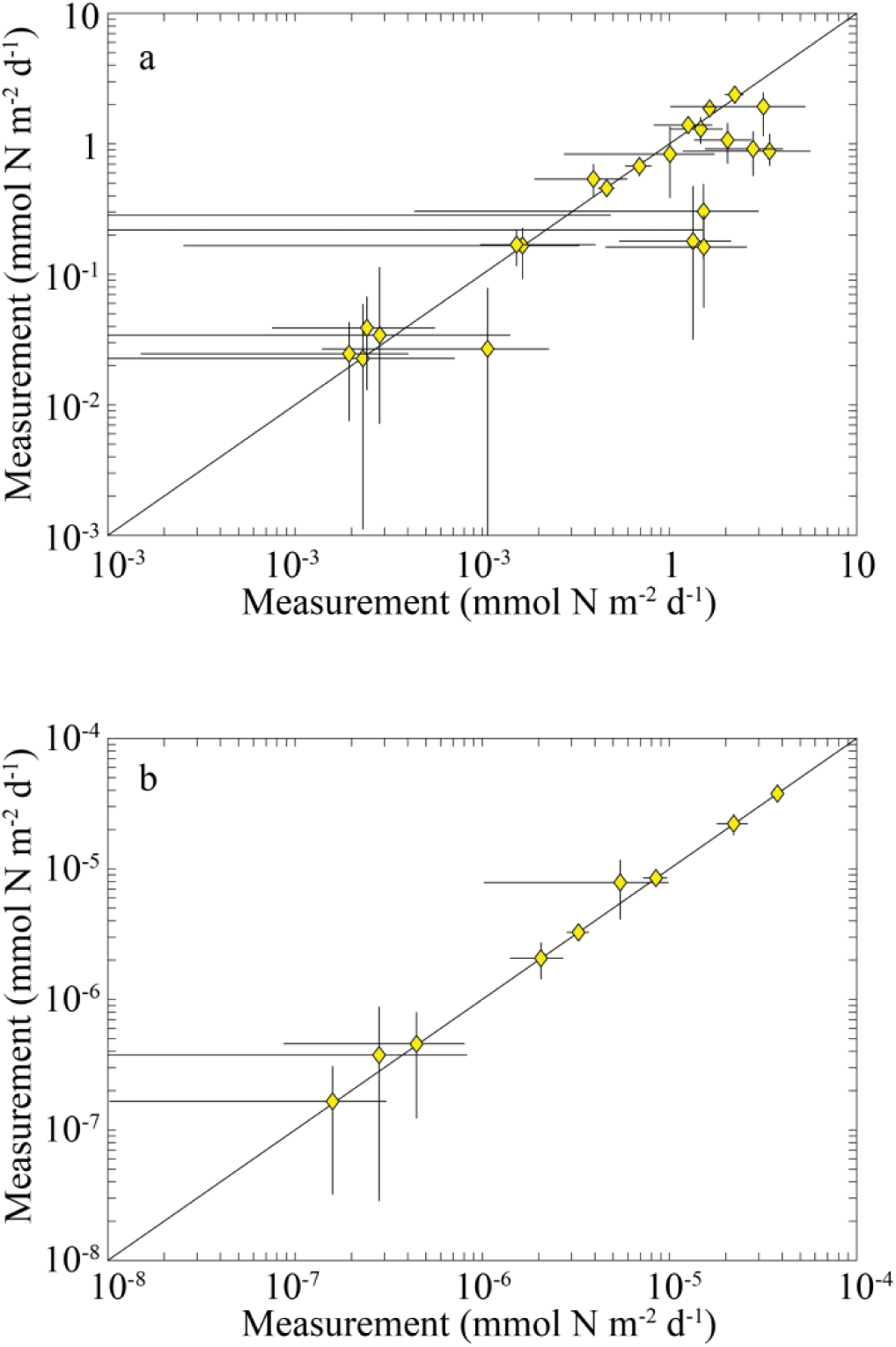
Comparison between field measurements and model estimates for planktonic ecosystem rate measurements (a) and ABT feeding measurements (b).

**Fig. 3.**
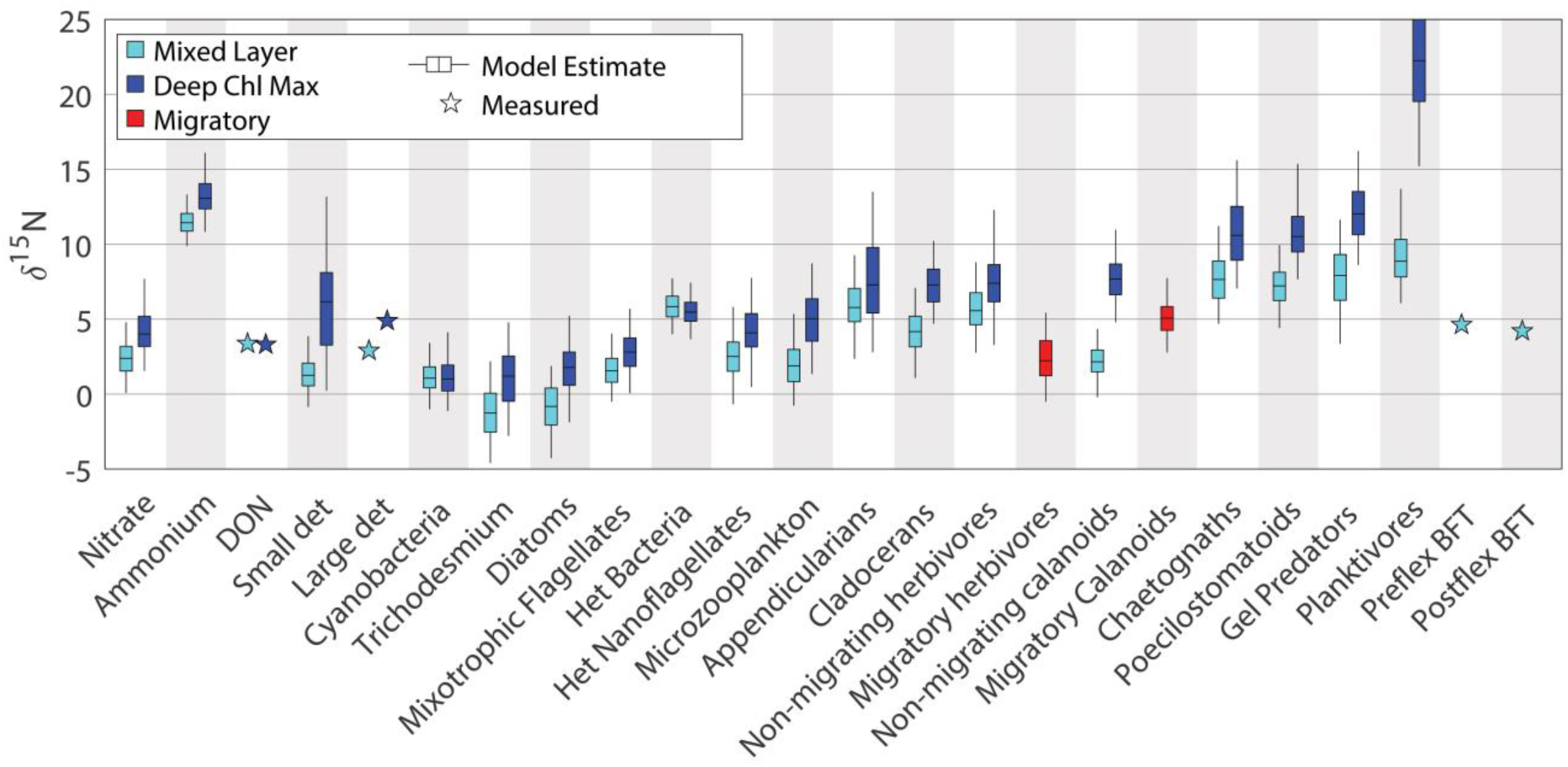
Boxplots (median, inter-quartile ranges, and 95% C.I.) for δ^15^N of mixed layer model compartments (light blue), deep euphotic zone compartments (dark blue), and vertically-migrating groups (red). Stars indicate field measurements.

### Food web dynamics

Food web dynamics broadly reflected those expected for an oligotrophic, recycling-dominant ecosystem. NH_4_^+^ was the dominant source of nitrogen to phytoplankton (mean = 82%; 95% C.I. = 70 – 92%). NO_3_^-^ uptake (15%; 6 – 27%) and N_2_ fixation (2.9%; 0.12 – 7.5%) were comparatively less important. Nutrient utilization patterns were broadly similar in the vicinity of the deep chlorophyll max (>50 m depth), although NH_4_^+^ uptake was slightly less important (59%; 27 – 88%) and NO_3_^-^ uptake was comparatively more important (40%; 11 – 72%), with a negligible role for N_2_ fixation (1%; 0.023 – 3.6%). Total productivity was slightly higher in the shallow euphotic zone (2.4 mmol N m^-2^ d^-1^; 2.2 – 2.6 mmol N m^-2^ d^-1^) than in the deep euphotic zone (1.9 mmol N m^-2^ d^-1^; 1.7 – 2.0 mmol N m^-2^ d^-1^).

Picophytoplankton (54%; 38% - 68%) and flagellates (39%; 25 – 53%) were responsible for most of the primary production in the shallow euphotic zone. These contributions were notably flipped relative to the field data, which suggested greater production by flagellates. The model-data discrepancy related to the overall over-estimation of taxon-specific NPP by the dilution method, relative to H^14^CO_3_^-^ uptake measurements, and also due to the fact that the solution vectors originating from cyanobacteria were more likely than those originating from flagellates, given other constraints on the ecosystem. Diatoms were comparatively less important (6.9%; 3.7 – 9.5%), while *Trichodesmium* production was negligible. The relative proportions of each group were similar at the deep chlorophyll max.

Phytoplankton mortality was dominated by protistan grazing. These zooplankton (including mixotrophic flagellates) consumed 67% (48 – 81%) of phytoplankton production. Metazoan zooplankton consumed a much lower portion of phytoplankton production (13%; 8 – 19%), although they consumed more of the production of diatoms than protists did. Suspension-feeding metazoans also relied heavily on protistan zooplankton as dietary sources. This was reflected in trophic positions that averaged greater than 3.0 for all metazoans except appendicularians (Fig. 4). Predatory zooplankton (poecilostomatoid copepods, chaetognaths, and gelatinous predators) had particularly high trophic positions of 4.3, 4.3, and 4.6, respectively, in the upper euphotic zone. Their mean trophic positions in the deep euphotic zone were slightly lower (4.0, 3.9, and 4.3, for poecilostomatoid copepods, chaetognaths, and gelatinous predators, respectively).

**Fig. 4.**
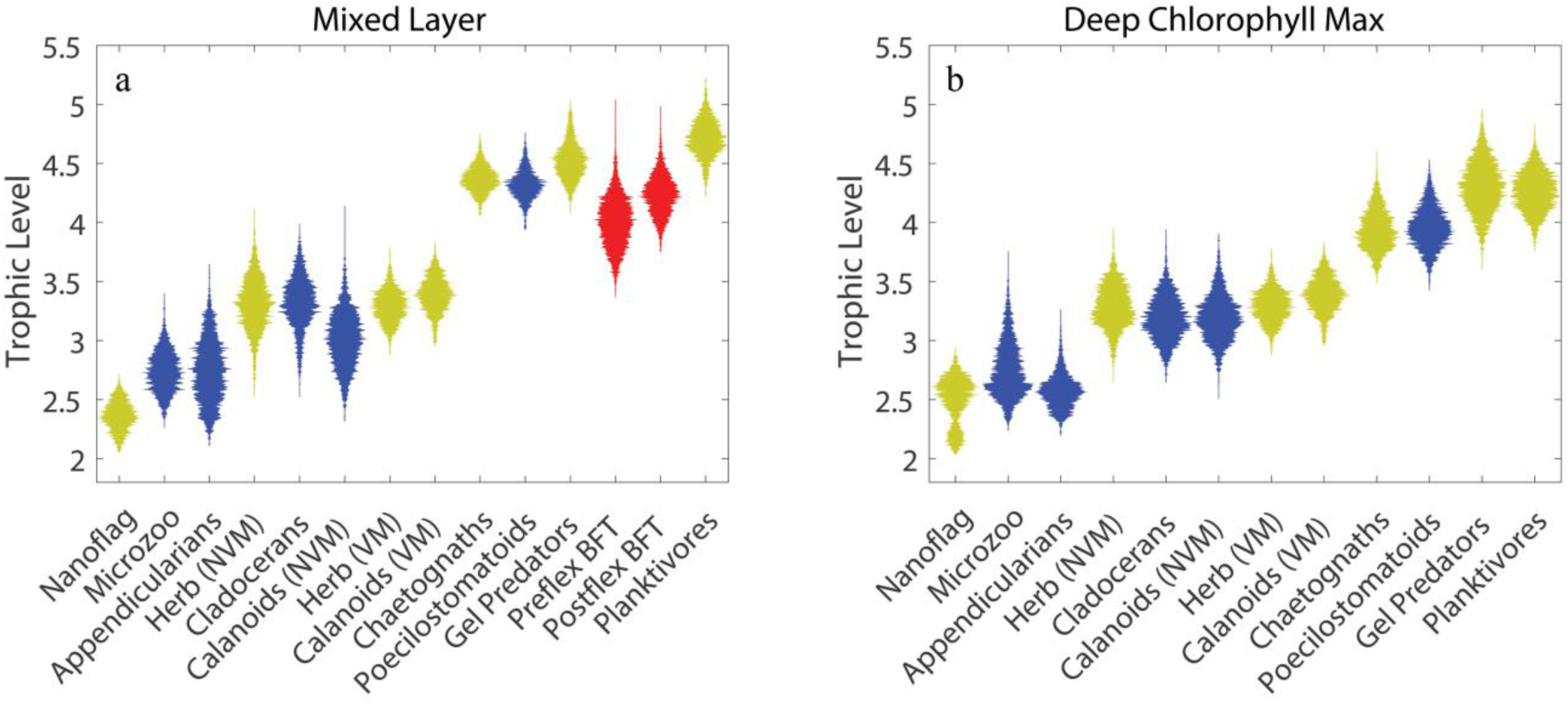
Violin plots of trophic level of zooplankton and fish in the mixed layer (a) and deep chl max (b). Blue plots are ABT prey. Red plots are ABT.

Quantification of major food web pathways showed that the GoM euphotic zone is dominated by the microbial loop (Fig. 5). The microbial loop (defined as the respiration of heterotrophic bacteria and the proportion of protistan respiration supported by bacterial production) processed 76% (52 – 106%) of net primary production in the shallow euphotic zone and 81% (68 – 92%) of net primary production in the deep euphotic zone. Note that microbial loop respiration can exceed net primary production, because net primary production does not include phytoplankton exudation of organic matter. For comparison, the herbivorous and multivorous food chains were responsible for 3.3% and 15% of NPP, respectively, in the shallow euphotic zone and 26% and 36%, respectively in the deep euphotic zone. The dominance of microbial loop pathways aligns with the importance of recycled NH_4_^+^ for phytoplankton production and conforms with an expectation of tight recycling in oligotrophic ecosystems with limited new nutrient supply. In the shallow euphotic zone, where recycling and the microbial loop were most important, phytoplankton and protistan zooplankton had approximately equal roles in DON production (32% and 33%, respectively), with much of the remainder coming from dissolution of detritus. Mesozooplankton excretion was responsible for only 3.3% of DOM production. Bacterial excretion was in turn responsible for 57% of NH_4_^+^ regeneration in the shallow euphotic zone, with protist excretion generating an additional 37% of the NH_4_^+^ used by phytoplankton.

**Fig. 5.**
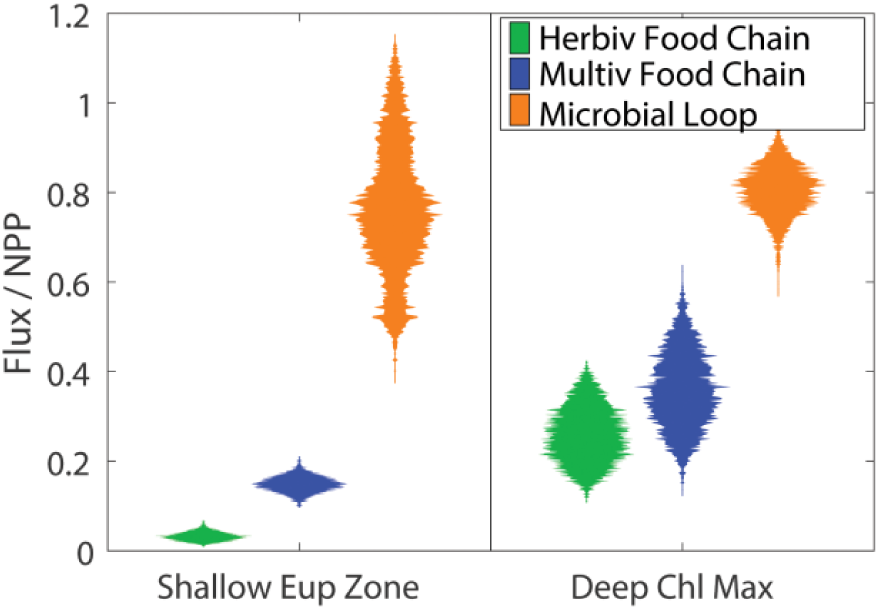
Flux through the herbivorous food chain, multivorous food chain, and microbial loop normalized to net primary production.

### Larval Atlantic Bluefin Tuna in the GoM Ecosystem

As suggested by the gut content data, our model results show that larval ABT feed predominantly on calanoid copepods, with a lesser role for microzooplankton, appendicularians, cladocerans, and poecilostomatoid copepods in their diets (Fig. 6). Calanoid copepods comprised 88.1% of the diet of preflexion ABT (95% C.I. = 77.4 – 94.9%). Microzooplankton (2.0%; C.I. = 0.4% - 4.2%), appendicularians (5.4%; 1.4 – 10.7%), and cladocerans (4.4%; 0.3 – 11.3%) were all small contributors to the diets of preflexion ABT, while poecilostomatoid copepods were negligible contributors to preflexion ABT diets (<0.3%). Although calanoid copepods were also the dominant dietary source for postflexion ABT (51.2%; 48.1 – 54.5%), these larger larvae had more varied diets with important contributions from cladocerans (30%; 25.9 – 33.9%) and poecilostomatoid copepods (11.5%; 9.9% - 13.1%).

**Fig. 6.**
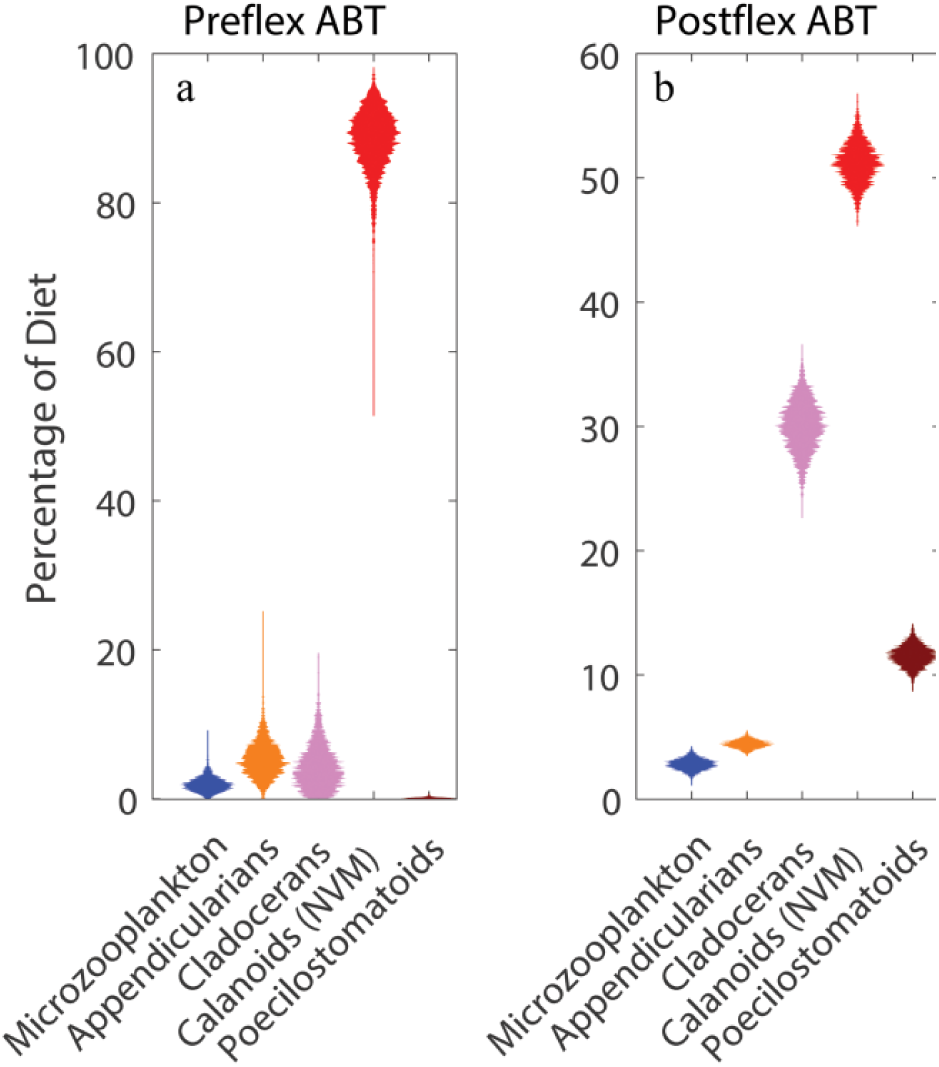
Violin plots of larval ABT diets

The prevalence of suspension-feeding zooplankton in the diets of both pre-flexion and post-flexion diets led to relatively low trophic levels for ABT larvae (Fig. 4). Given the ecosystem structure used in the model (Fig. 1), larval ABT could potentially have a trophic level between 3 and 7. However, both preflexion and postflexion larvae had trophic levels on the low end of this range. Preflexion ABT had a trophic level of 4 (95% C.I. = 3.6 – 4.5), while postflexion ABT had a trophic level of 4.2 (3.9 −4.5). Both groups of ABT larvae thus had a trophic level that averaged approximately 0.6 of their maximum possible trophic level (Fig. 7) and only one trophic level higher than the lowest theoretically possible trophic level they could take on within the food web. The trophic positions of larval ABT were thus notably low relative to the trophic positions they would take on if feeding on the longest possible food chains that the model allowed. Based on this metric, they were also notably lower in trophic position than many of the zooplankton and other fish in the model.

**Fig. 7.**
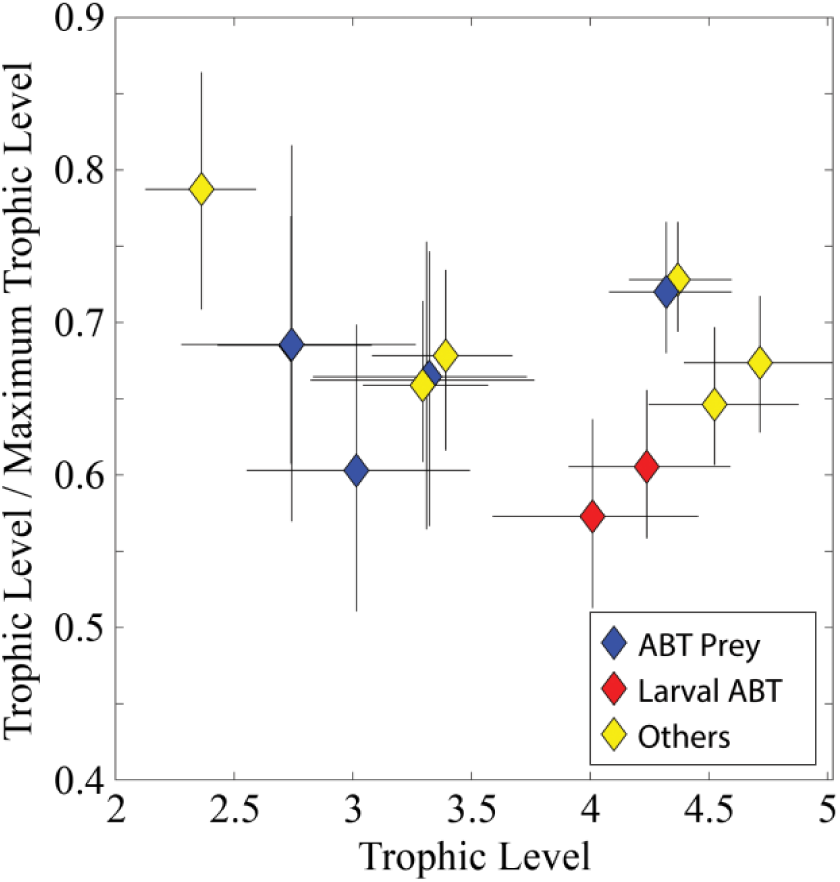
The ratio of the trophic level of different modeled zooplankton and fish to the trophic level they would have in the longest possible model food chain.

The food chains supporting larval ABT were diverse and relied on significant production of picophytoplankton, flagellates, and diatoms (while the production of *Trichodesmium* was insignificant for ABT food chains). Preflexion ABT excreted 1.6 nmol N m^-2^ d^-1^ (0.58 – 3.4 nmol N m^-1^ d^-1^) derived from the production of flagellates, 0.97 (0.34 – 2.1) nmol N m^-1^ d^-1^ from picophytoplankton, and 0.75 (0.17 – 1.7) nmol N m^-1^ d^-1^ from diatoms (Fig. 8a). Post-flexion larvae excreted 12 (6.4 – 23) nmol N m^-1^ d^-1^ from flagellates, 7.6 (3.5 – 15) nmol N m^-1^ d^-1^ from picophytoplankton, and 4.3 (1.5 – 8.6) nmol N m^-1^ d^-1^. These values were influenced in large part by the different production rates of each phytoplankton taxa (flagellate, picophytoplankton, and diatom NPP in the shallow euphotic zone were 2.0, 1.5, and 0.1 mmol N m^-2^ d^-1^). When normalized to phytoplankton NPP it becomes clear that larval tuna rely disproportionately on the production of large phytoplankton (Fig. 8b). Pre-flexion ABT respired 3.6×10^−4^ % of diatom NPP and 1.1×10^−4^ % of flagellate NPP, compared to only 5.7×10^−5^ % of picophytoplankton NPP. Post-flexion larvae respired 2.1×10^−3^ % of diatom NPP, 8.4×10^−4^ % of flagellate NPP, and 4.4×10^−4^ % of picophytoplankton NPP. The proportion of *Trichodesmium* NPP respired by larvae was poorly constrained by the model, although *Trichodesmium* production was consistently low in all model solution vectors. The disproportionately large role of diatoms in larval ABT diets was reflected in the roles of diatoms in supporting their mesozooplankton prey (Fig. 8d). All four mesozooplankton prey taxa respired a higher proportion of diatom taxa than any other phytoplankton and three of the four had flagellates as the second most important phytoplankton taxa. Only appendicularians excreted a higher proportion of picophytoplankton production than flagellate production. These results for mesozooplankton were in stark contrast to similar proportional roles for phytoplankton in protists (Fig. 8c). Heterotrophic nanoflagellates relied disproportionately on picophytoplankton, respiring 26% of picophytoplankton NPP, while microzooplankton relied disproportionately on the NPP of flagellates (respiring 29% of flagellate NPP).

**Fig. 8.**
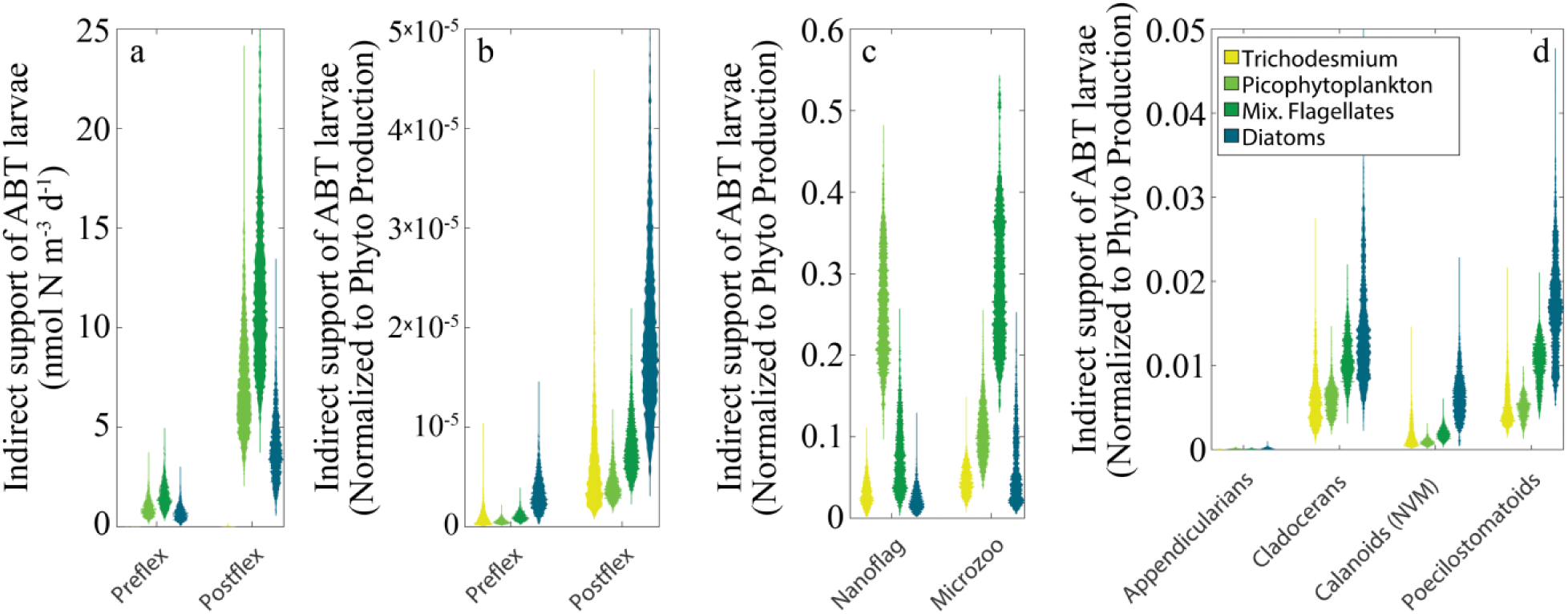
Indirect foodweb flows to larval tuna (a, b), protists (c), and mesozooplankton (d). Panel a shows the amount of organic matter derived from each phytoplankton taxon that was respired by larval tuna. Panels b, c, and d show the proportion of the production of each phytoplankton taxon that was respired by either larval tuna (b), protists (c), or mesozooplankton (d). Only ABT prey are shown in d.

## DISCUSSION

The open-ocean Gulf of Mexico is an incredibly oligotrophic ecosystem with low productivity and a deep nitracline (Biggs, 1992, Gomez *et al.*, 2018, Knapp *et al.*, in prep., Yingling *et al.*, in prep.). Nevertheless, it is an important spawning ground for many migratory fish species, including multiple species of tuna, dolphinfish, sailfish, and marlin (Cornic *et al.*, 2018, Kitchens & Rooker, 2014, Rooker *et al.*, 2012). Models suggest that vertical mixing is an important structuring process for productivity in the open-ocean Gulf (Gomez *et al.*, 2018). If vertical mixing is indeed crucial for supporting these oligotrophic systems, predicted future warming and increased stratification could have deleterious impacts on taxa living in the mixed layer (Liu *et al.*, 2015, Muhling *et al.*, 2011, Muhling *et al.*, 2015). Understanding how pelagic ecosystems, and the larval fish they support, will respond to climate change requires knowledge of the food web pathways that convert phytoplankton production into the preferred prey of different species (Landry *et al.*, 2019).

We can hypothesize two potential ways in which an organism’s diet could make it well adapted to life in an oligotrophic region. First, it could feed preferentially on taxa that have either direct or indirect linkages to some of the most abundant primary producers in the ecosystem (e.g., cyanobacteria). For instance, a reliance on appendicularians would give larval fish access to a suspension-feeder that can feed directly on picophytoplankton (Gorsky & Fenaux, 1998, Llopiz *et al.*, 2010). Conversely, preference for calanoid copepods would make a larval fish more dependent on the production of diatoms and other large phytoplankton. A second, but not mutually-exclusive, hypothesis states that larval fish are more likely to thrive in oligotrophic ecosystems if they feed at a low trophic position, thus maximizing trophic transfer efficiency from phytoplankton to larvae, regardless of the source of the production.

Our results show no evidence for the former hypothesis. Although diatom production only contributed to ∼20% of ABT larval diets, this was a disproportionately high fraction of the ABT diet relative to the proportional role of diatoms to total primary production (<10%). Indeed, relative to a phytoplankton taxon’s productivity, the proportional contribution of each phytoplankton taxon to foodweb pathways that support ABT larvae increased with increasing phytoplankter size from picophytoplankton to flagellates to diatoms (Fig. 8). The disproportionately large role of diatom-driven pathways was largely the result of non-vertically migrating calanoid copepods (and to a lesser extent cladocerans), which formed a large portion of the diet of ABT larvae and fed substantially on diatoms. In contrast, while an efficient pathway from cyanobacteria to ABT larvae existed (through appendicularians), appendicularians were not abundant in either ABT guts or in the water column. Instead, the majority of cyanobacteria were consumed by heterotrophic nanoflagellates. These heterotrophic nanoflagellates had a relatively low gross growth efficiency in the model (∼20%) and their primary predators were other protists (microzooplankton). Cyanobacteria and heterotrophic nanoflagellates thus contributed substantially to the recycling pathways of the microbial loop, forming a largely distinct food web from the multivorous and herbivorous pathways, which mostly began with mixotrophic flagellates and diatoms and supported the production of larval ABT and other planktivorous fish.

Our results offer more support for the hypothesis that ABT larvae feed at a relatively low trophic level to maximize the proportion of primary productivity available to them (Fig. 7). The trophic position of ABT larvae (∼4) is much closer to the minimum trophic level that our model allows (3: phytoplankton→prey→larvae) than to the maximum allowed trophic level (7: phytoplankton→bacteria→nanoflagellates→microzooplankton→suspension-feeders →carnivorous zooplankton→larvae). The low trophic position of ABT larvae is particularly striking considering how weak the herbivorous food chain is. Generally, planktivorous fish are more likely to be at a low trophic level if they are feeding on an ecosystem dominated by large phytoplankton and herbivorous mesozooplankton. However, the herbivorous food chain was responsible for only 3.3% of net primary production in the shallow euphotic zone where ABT larvae feed; the multivorous food chain processed 15% of NPP, while the microbial loop processed 76% (Fig. 5). The low trophic position of ABT larvae was primarily due to two factors: 1) although total protistan secondary production was higher than total mesozooplankton secondary production, a comparatively small proportion of this secondary production proceeded to food chains ending in larval tuna. Instead, much of it was dissipated as respiration within the microbial loop. Food chains supporting larval ABT were largely distinct from food chains involving the smallest class of heterotrophic protists. 2) Both size classes of ABT larvae fed primarily on non-vertically migrating calanoid copepods and these calanoid copepods tended to feed lower on the food chain than other similar suspension-feeding taxa.

While this trophic level of ∼4 is low for a species known to preferentially feed on carnivorous copepods (poecilostomatoids) in a cyanobacteria and microbial loop-driven ecosystem, it is important to note that this is not actually a low trophic level relative to some other mass-balance constrained marine food web models. Many models based on ECOPATH software include only one (or zero) protistan trophic steps and many include a single mesozooplankton group (Arreguin-Sanchez *et al.*, 2004, Geers *et al.*, 2016, Walters *et al.*, 2008). Thus many of these models constrain zooplankton to a trophic level of 2 or 3 and hence the maximum allowed trophic level for planktivores becomes only 3 or 4. The additional complexity associated with planktonic ecosystems in our model is a far more realistic depiction of the complexity of these food webs (Fig. 1). Nevertheless, our model only allows a maximum of two trophic steps within the protistan zooplankton (heterotrophic nanoflagellates and microzooplankton). This is in many ways an artificial distinction. Protistan food webs are fluid. Some protists (e.g., pallium-feeding dinoflagellates) routinely feed at a 1:1 predator:prey size ratio, while others (e.g., ciliates) often feed at closer to a 10:1 predator:prey size ratio (Fuchs & Franks, 2010, Kiorboe, 2008). Some protists are thus likely to take on a higher trophic position than the maximum value allowed by our model. Indeed, the high energy dissipation rates that our model finds for protists (model average for heterotrophic nanoflagellate gross growth efficiency is only ∼20%) suggest that other model constraints (e.g., δ^15^N of consumers and sinking detritus) are pushing the model towards lower trophic efficiency amongst the protistan food web, possibly indicating that there are in fact more trophic links amongst this group than we allow.

Although our cruise plan was not designed to focus on spatial variability and mesoscale features, it is important to note that mesoscale eddies may play an important role in structuring GoM ecosystems. The GoM is a region with very high mesoscale energy, with the Loop Current (and the eddies that it sheds) forming some of the most prominent (but certainly not only) features in the open-ocean GoM (Forristall *et al.*, 1992, Oey *et al.*, 2005, Schmitz *et al.*, 2005). These features have the potential to fundamentally restructure open-ocean ecosystems, with warm-core eddies (including Loop Current Eddies) depressing the nutricline and hence primary production and cold-core eddies leading to increased open-ocean upwelling and productivity (Biggs & Müller-Karger, 1994). These altered nutrient supply and phytoplankton regimes lead to substantially higher zooplankton biomass within cold-core eddies (Wells *et al.*, 2017). However, the relative importance of each eddy type, as well as the distinct gradient regions that form on their edges, on larval ABT remains a topic of active debate (Domingues *et al.*, 2016, Muhling *et al.*, 2010, Shropshire *et al.*, in prep.). Determining the response of pelagic food webs and ABT larval tuna to climate change will thus require characterizing not only how GoM circulation will respond to a future climate, but also food web variability across these features (Liu *et al.*, 2015, Muhling *et al.*, 2011). Our study offers substantial insight into the processes allowing larval ABT to survive in a food-scare environment. However, substantial future research is necessary to quantify the impact of spatial and interannual variability, as well as secular change, on these ecosystems and potentially threatened species.

## CONCLUSIONS

The larvae of ABT feed in oligotrophic ecosystems, dominated by cyanobacteria and other small phytoplankton. These ecosystems are dominated by the microbial loop, within which both bacteria and protists play important roles in regenerating nutrients. However, larval ABT feed preferentially on less dominant food web pathways associated with herbivorous and multivorous food chains. Consequently, they are reliant on the production of diatoms and mixotrophic flagellates that support herbivorous zooplankton taxa, particularly calanoid copepods and cladocerans. Preferential utilization of these more direct trophic pathways allows larval ABT to feed at relatively low trophic levels despite the fact that the taxa responsible for the majority of secondary production in the food web (bacteria and heterotrophic nanoflagellates) are not accessible as prey to them. Further research is needed to understand how these ecological interactions shift under different disturbance regimes.

## ACKNOWLEDGMENTS

We thank our numerous colleagues in the BLOOFINZ-GoM project who made this research possible. Our work was funded by National Science Foundation Biological Oceanography grant #1851347 and a National Oceanic and Atmospheric Administration’s RESTORE Program Grant (Project Title: Effects of nitrogen sources and plankton food-web dynamics on habitat quality for the larvae of Atlantic bluefin tuna in the Gulf of Mexico) under federal funding opportunity NOAA-NOS-NCCOS-2017-2004875. https://restoreactscienceprogram.noaa.gov/funded-projects/bluefin-tuna-larvae. Data utilized in this manuscript are available on NCCOS and BCO-DMO (https://www.bco-dmo.org/project/819488).

---

**Supplementary Table 1.**
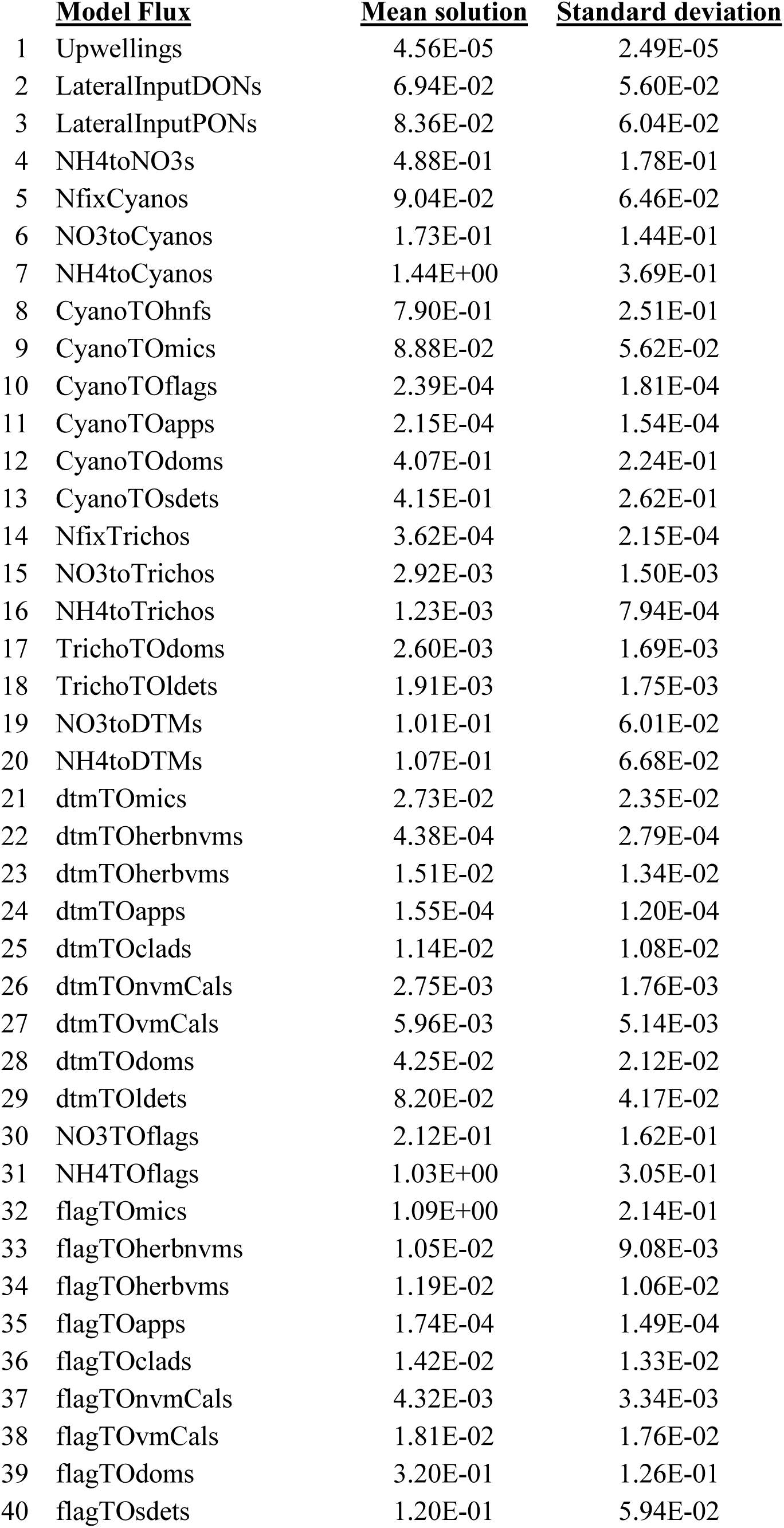
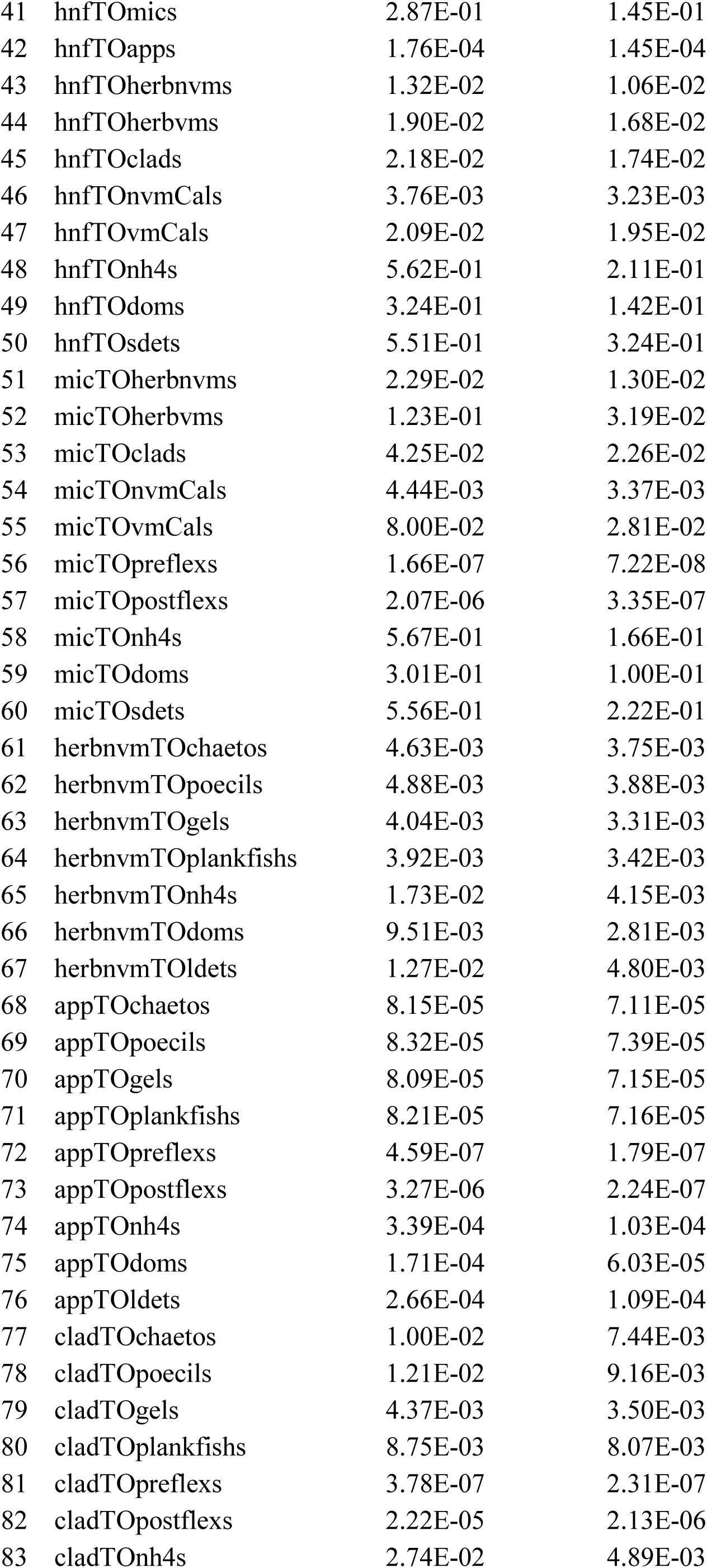
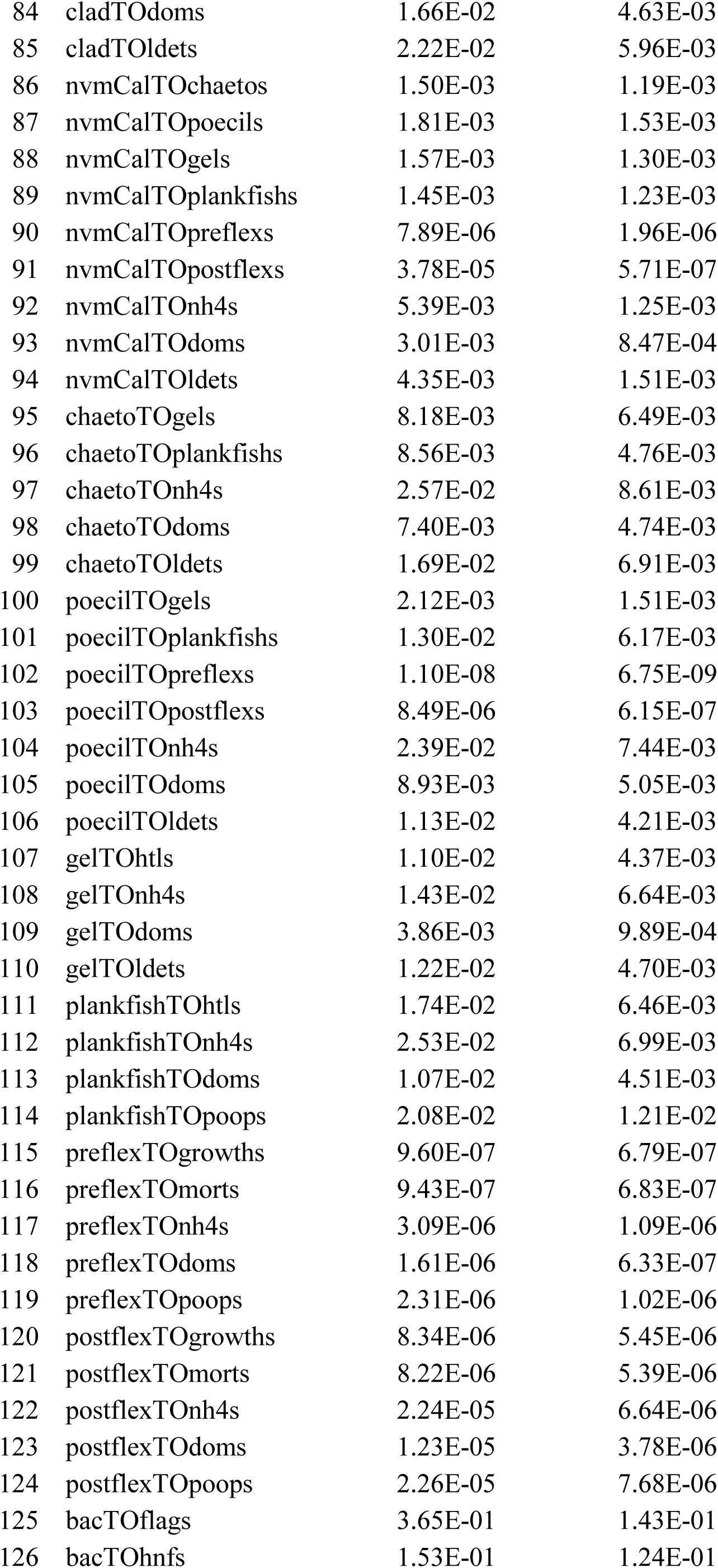
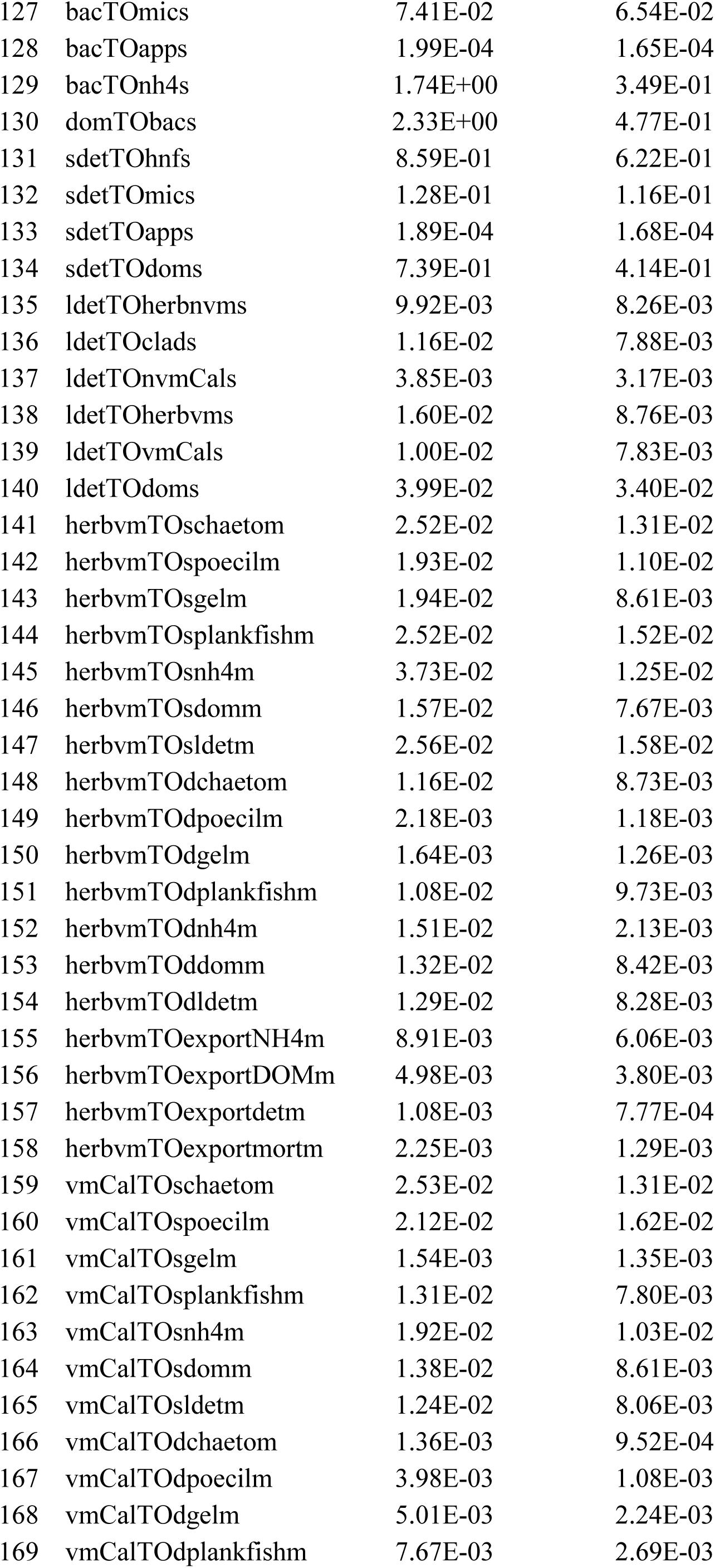
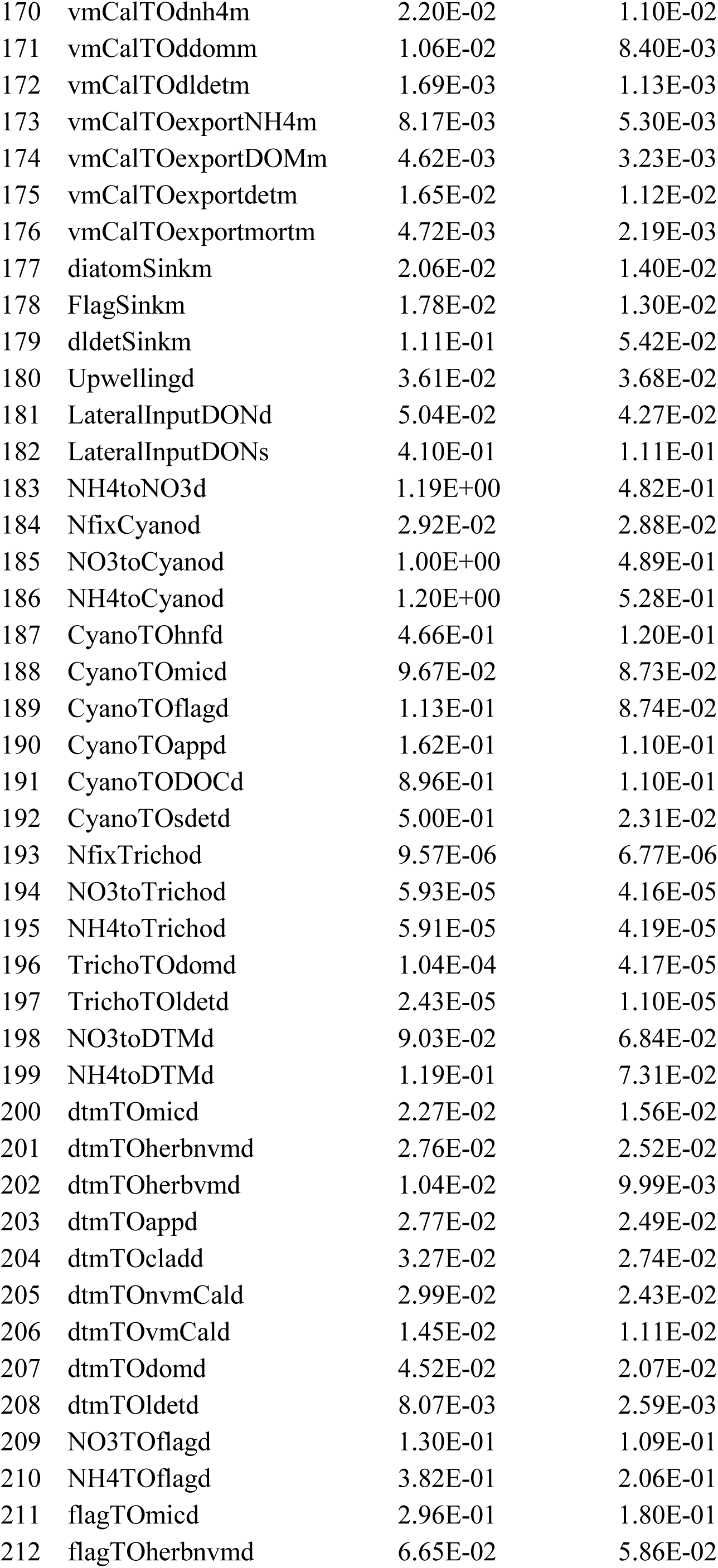
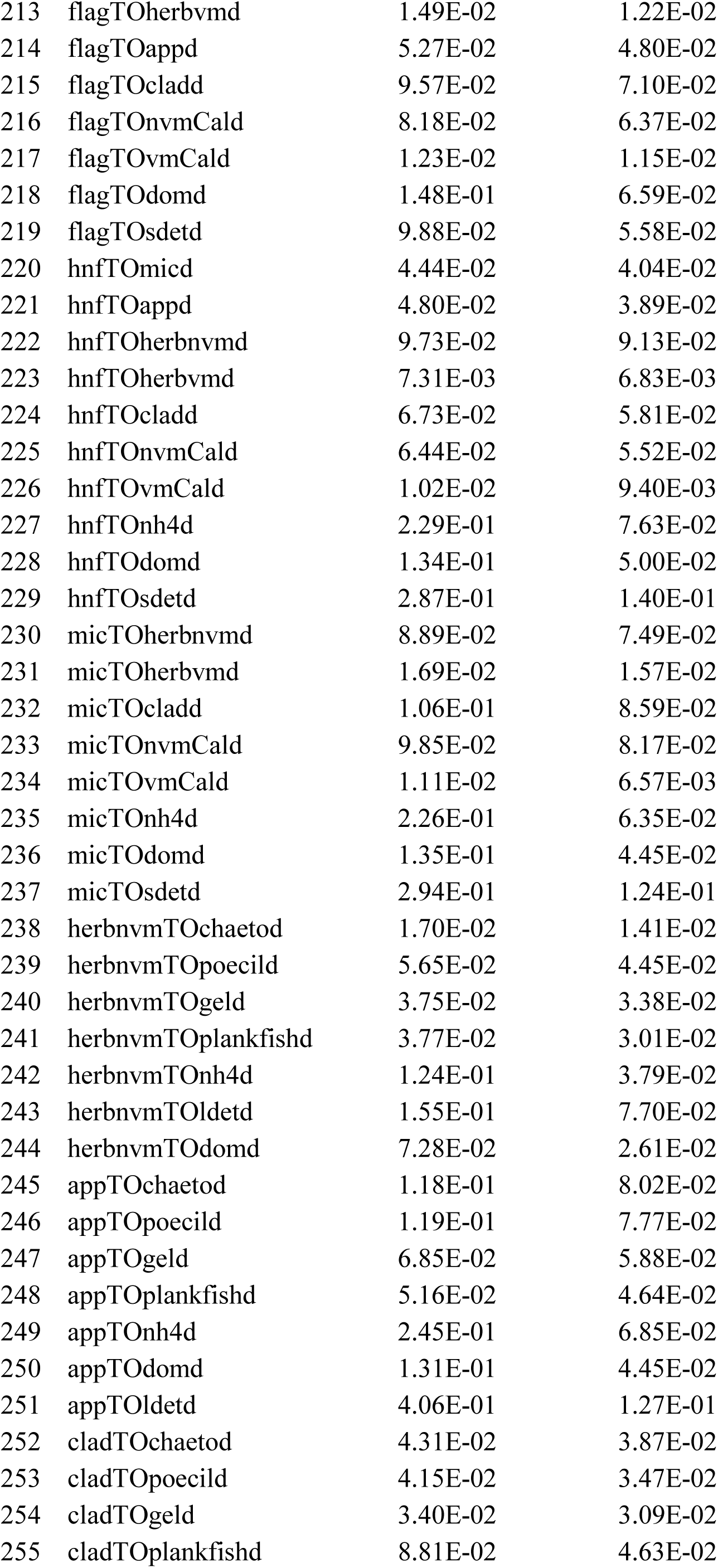
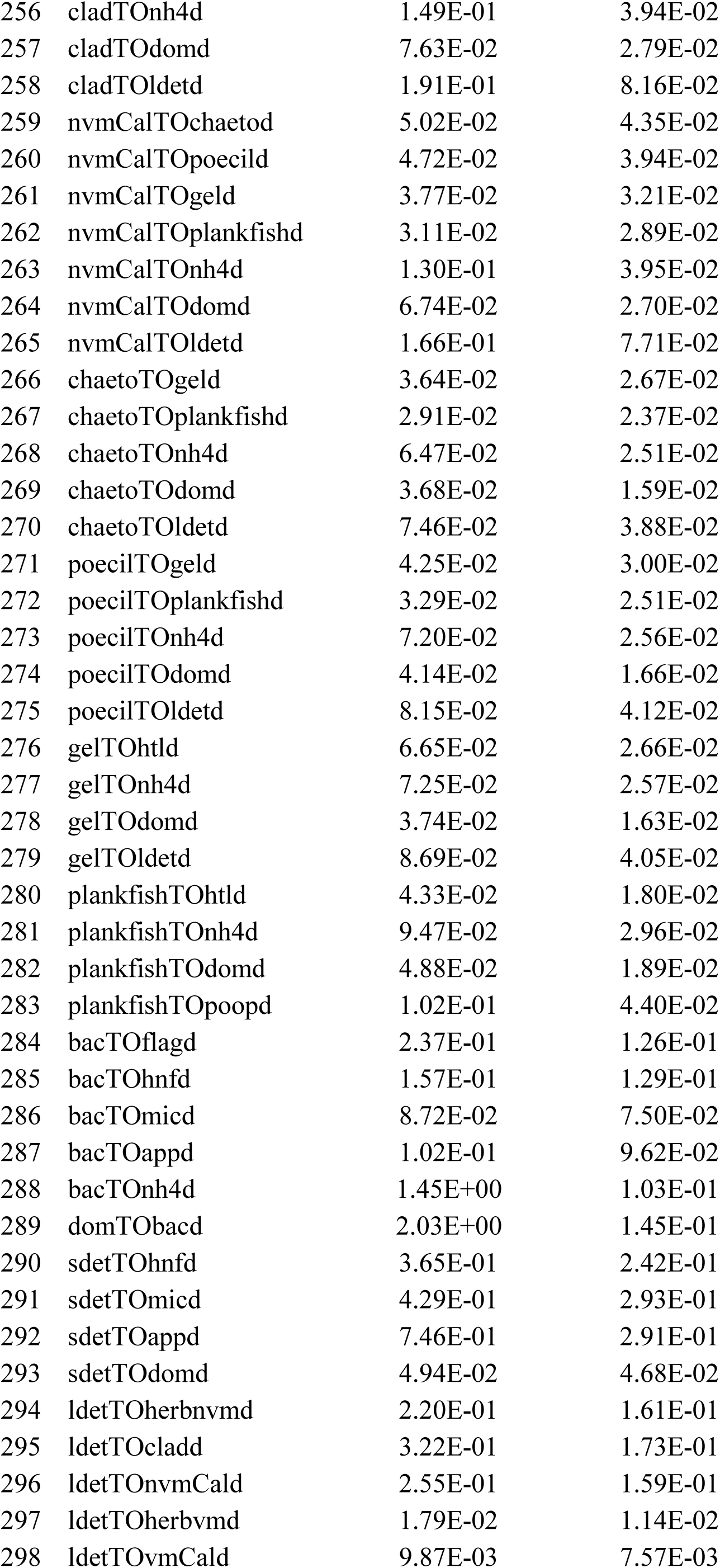
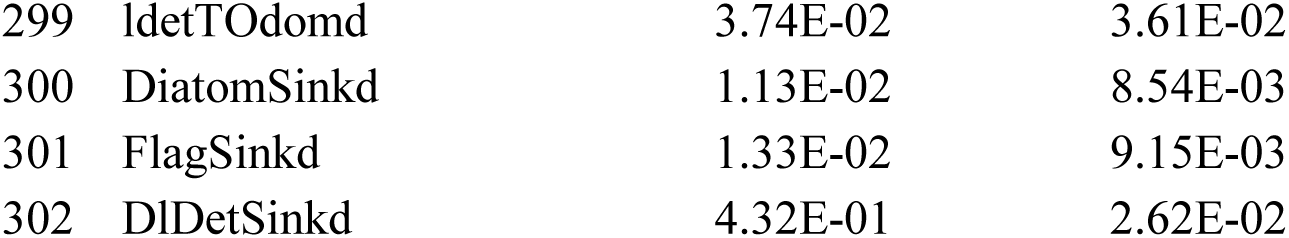
Model Solutions.

## REFERENCES

Alldredge, A. (1976) Field behavior and adaptive strategies of appendicularians (Chordata: Tunicata). Marine Biology, 38, 29–39.

Arreguin-Sanchez, F., Zetina-Rejón, M., Manickchand-Heileman, S., Ramirez-Rodriguez, M. and Vidal, L. (2004) Simulated response to harvesting strategies in an exploited ecosystem in the southwestern Gulf of Mexico. ecological modelling, 172, 421–432.

Barton, A. D., Irwin, A. J., Finkel, Z. V. and Stock, C. A. (2016) Anthropogenic climate change drives shift and shuffle in North Atlantic phytoplankton communities. Proceedings of the National Academy of Sciences, 113, 2964–2969.

Biggs, D. C. (1992) Nutrients, plankton, and productivity in a warm-core ring in the western Gulf of Mexico. Journal of Geophysical Research: Oceans, 97, 2143–2154.

Biggs, D. C. and Müller-Karger, F. E. (1994) Ship and satellite observations of chlorophyll stocks in interacting cyclone-anticyclone eddy pairs in the western Gulf of Mexico. Journal of Geophysical Research: Oceans, 99, 7371–7384.

Biggs, D. C. and Ressler, P. H. (2001) Distribution and abundance of phytoplankton, zooplankton, ichthyoplankton, and micronekton in the deepwater Gulf of Mexico. Gulf of Mexico Science, 19, 2.

Bode, M., Hagen, W., Schukat, A., Teuber, L., Fonseca-Batista, D., Dehairs, F. and Auel, H. (2015) Feeding strategies of tropical and subtropical calanoid copepods throughout the eastern Atlantic Ocean - Latitudinal and bathymetric aspects. Progress in Oceanography, 138, 268–282.

Cornic, M., Smith, B. L., Kitchens, L. L., Bremer, J. R. A. and Rooker, J. R. (2018) Abundance and habitat associations of tuna larvae in the surface water of the Gulf of Mexico. Hydrobiologia, 806, 29–46.

Damien, P., Pasqueron De Fommervault, O., Sheinbaum, J., Jouanno, J., Camacho-Ibar, V. F. and Duteil, O. (2018) Partitioning of the Open Waters of the Gulf of Mexico Based on the Seasonal and Interannual Variability of Chlorophyll Concentration. Journal of Geophysical Research: Oceans, 123, 2592–2614.

Décima, M., Landry, M. R., Stukel, M. R., Lopez-Lopez, L. and Krause, J. W. (2016) Mesozooplankton biomass and grazing in the Costa Rica Dome: amplifying variability through the plankton food web. Journal of Plankton Research, 38, 317–330.

Décima, M., Stukel, M. R., López-López, L. and Landry, M. R. (2019) The unique ecological role of pyrosomes in the Eastern Tropical Pacific. Limnology and Oceanography, 64, 728–743.

Domingues, R., Goni, G., Bringas, F., Muhling, B., Lindo-Atichati, D. and Walter, J. (2016) Variability of preferred environmental conditions for Atlantic bluefin tuna (Thunnus thynnus) larvae in the Gulf of Mexico during 1993–2011. Fisheries Oceanography, 25, 320–336.

Flombaum, P., Gallegos, J. L., Gordillo, R. A., Rincón, J., Zabala, L. L., Jiao, N., Karl, D. M., Li, W. K. W., Lomas, M. W., Veneziano, D., Vera, C. S., Vrugt, J. A. and Martiny, A. C. (2013) Present and future global distributions of the marine Cyanobacteria Prochlorococcus and Synechococcus. Proceedings of the National Academy of Sciences, 110, 9824–9829.

Forristall, G. Z., Schaudt, K. J. and Cooper, C. K. (1992) Evolution and kinematics of a Loop Current eddy in the Gulf of Mexico during 1985. Journal of Geophysical Research: Oceans, 97, 2173–2184.

Fuchs, H. L. and Franks, P. J. S. (2010) Plankton community properties determined by nutrients and size-selective feeding. Marine Ecology Progress Series, 413, 1–15.

Gargett, A. and Garner, T. (2008) Determining Thorpe scales from ship-lowered CTD density profiles. Journal of Atmospheric and Oceanic Technology, 25, 1657–1670.

Geers, T. M., Pikitch, E. K. and Frisk, M. G. (2016) An original model of the northern Gulf of Mexico using Ecopath with Ecosim and its implications for the effects of fishing on ecosystem structure and maturity. Deep Sea Research Part II: Topical Studies in Oceanography, 129, 319–331.

Gomez, F. A., Lee, S.-K., Liu, Y., Hernandez Jr, F. J., Muller-Karger, F. E. and Lamkin, J. T. (2018) Seasonal patterns in phytoplankton biomass across the northern and deep Gulf of Mexico: a numerical model study. Biogeosciences, 15, 3561–3576.

Gorsky, G. and Fenaux, R. (1998) The role of Appendicularia in marine food webs. The biology of pelagic tunicates, 161–169.

Hannon, B. (1973) Structure of ecosystems. Journal of Theoretical Biology, 41, 535–546.

Hong, H., Shen, R., Zhang, F., Wen, Z., Chang, S., Lin, W., Kranz, S. A., Luo, Y.-W., Kao, S.-J. and Morel, F. M. (2017) The complex effects of ocean acidification on the prominent N2-fixing cyanobacterium Trichodesmium. Science, 356, 527–531.

Ikeda, T. (1985) Metabolic rates of epipelagic marine zooplankton as a function of body mass and temperature. Marine Biology, 85, 1–11.

Katechakis, A. and Stibor, H. (2004) Feeding selectivities of the marine cladocerans Penilia avirostris, Podon intermedius and Evadne nordmanni. Marine Biology, 145, 529–539.

Kelly, T. B., Landry, M. R., Selph, K. E., Knapp, A. N., Swalethorp, R. and Stukel, M. R. (in prep.) The balance of new and export production in the oligotrophic Gulf of Mexico. Journal of Plankton Research.

Kiorboe, T. (2008) A Mechanistic Approach to Plankton Ecology. Vol., Princeton University Press, Princeton, NJ.

Kitchens, L. L. and Rooker, J. R. (2014) Habitat associations of dolphinfish larvae in the Gulf of Mexico. Fisheries Oceanography, 23, 460–471.

Knapp, A. N., Sigman, D. M. and Lipschultz, F. (2005) N isotopic composition of dissolved organic nitrogen and nitrate at the Bermuda Atlantic time-series study site. Global Biogeochemical Cycles, 19.

Knapp, A. N., Thomas, R., Stukel, M. R., Kelly, T. B., Landry, M. R., Selph, K. E., Malca, E., Gerard, T. and Lamkin, J. (in prep.) Constraining the sources of nitrogen fueling phytoplankton and food webs in the Gulf of Mexico using nitrogen isotope budgets. Journal of Plankton Research.

Kones, J. K., Soetaert, K., Van Oevelen, D. and Owino, J. O. (2009) Are network indices robust indicators of food web functioning? A Monte Carlo approach. Ecological Modelling, 220, 370–382.

Laiz-Carrion, R., Gerard, T., Uriarte, A., Malca, E., Quintanilla, J. M., Muhling, B. A., Alemany, F., Privoznik, S. L., Shiroza, A., Lamkin, J. T. and Garcia, A. (2015) Trophic Ecology of Atlantic Bluefin Tuna (Thunnusthynnus) Larvae from the Gulf of Mexico and NW Mediterranean Spawning Grounds: A Comparative Stable Isotope Study. Plos One, 10.

Landry, M. R. (2009) Grazing processes and secondary production in the Arabian Sea: A simple food web synthesis with measurement constraints. In: J. D. Wiggert, R. R. Hood, S. W. A. Naqvi, K. H. Brink and S. L. Smith (eds) Indian Ocean biogeochemical processes and ecological variability AGU Monograph. American Geophysical Union, pp. 133–146.

Landry, M. R., Beckley, L. E. and Muhling, B. A. (2019) Climate sensitivities and uncertainties in food-web pathways supporting larval bluefin tuna in subtropical oligotrophic oceans. Ices Journal of Marine Science, 76, 359–369.

Landry, M. R. and Lamkin, J. (in prep.) Pelagic ecology of the oligotrophic Gulf of Mexico: Overview of the BLOOFINZ-GoM Cruises. Journal of Plankton Research.

Landry, M. R., Selph, K. E., Decima, M., Gutierrez-Rodriquez, A., Stukel, M. R., Taylor, A. G. and Pasulka, A. L. (2016) Phytoplankton production and grazing balances in the Costa Rica Dome. Journal of Plankton Research, 38, 366–379.

Landry, M. R., Selph, K. E., Stukel, M. R., Swalethorp, R., Kelly, T. B., Beatty, J. and Quackenbush, C. (in prep.) Microbial food web dynamics in the oceanic Gulf of Mexico. Journal of Plankton Research.

Lindo-Atichati, D., Bringas, F., Goni, G., Muhling, B., Muller-Karger, F. E. and Habtes, S. (2012) Varying mesoscale structures influence larval fish distribution in the northern Gulf of Mexico. Marine Ecology Progress Series, 463, 245–257.

Liu, Y., Lee, S.-K., Enfield, D. B., Muhling, B. A., Lamkin, J. T., Muller-Karger, F. E. and Roffer, M. A. (2015) Potential impact of climate change on the Intra-Americas Sea: Part-1. A dynamic downscaling of the CMIP5 model projections. Journal of Marine Systems, 148, 56–69.

Llopiz, J. K., Muhling, B. A. and Lamkin, J. T. (2015) Feeding dynamics of Atlantic Bluefin Tuna (Thunnus thynnus) larvae in the Gulf of Mexico. Collect Vol Sci Pap ICCAT, 71, 1710–1715.

Llopiz, J. K., Richardson, D. E., Shiroza, A., Smith, S. L. and Cowen, R. K. (2010) Distinctions in the diets and distributions of larval tunas and the important role of appendicularians. Limnology and Oceanography, 55, 983–996.

Malca, E., Muhling, B., Franks, J., García, A., Tilley, J., Gerard, T., Ingram Jr, W. and Lamkin, J. T. (2017) The first larval age and growth curve for bluefin tuna (Thunnus thynnus) from the Gulf of Mexico: comparisons to the Straits of Florida, and the Balearic Sea (Mediterranean). Fisheries Research, 190, 24–33.

Malca, E., Quintanilla, J. M., Gerard, T., Laiz-Carrión, R., Garcia, A. and Lamkin, J. (in prep.) Spatial variability in larval growth between two spawning grounds: calibration and isotope analysis Journal of Plankton Research.

Mauchline, J., Blaxter, J. H. S., Southward, A. J. and Tyler, P. A. (1998) Advances in marine biology - The biology of calanoid copepods - Introduction Advances in Marine Biology, Vol 33. Vol. 33. pp. 1-+.

Muhling, B. A., Lamkin, J. T. and Roffer, M. A. (2010) Predicting the occurrence of Atlantic bluefin tuna (Thunnus thynnus) larvae in the northern Gulf of Mexico: building a classification model from archival data. Fisheries Oceanography, 19, 526–539.

Muhling, B. A., Lee, S.-K., Lamkin, J. T. and Liu, Y. (2011) Predicting the effects of climate change on bluefin tuna (Thunnus thynnus) spawning habitat in the Gulf of Mexico. Ices Journal of Marine Science, 68, 1051–1062.

Muhling, B. A., Liu, Y., Lee, S.-K., Lamkin, J. T., Roffer, M. A., Muller-Karger, F. and Walter Iii, J. F. (2015) Potential impact of climate change on the Intra-Americas Sea: Part 2. Implications for Atlantic bluefin tuna and skipjack tuna adult and larval habitats. Journal of Marine Systems, 148, 1–13.

Muller-Karger, F. E., Smith, J. P., Werner, S., Chen, R., Roffer, M., Liu, Y., Muhling, B., Lindo-Atichati, D., Lamkin, J. and Cerdeira-Estrada, S. (2015) Natural variability of surface oceanographic conditions in the offshore Gulf of Mexico. Progress in Oceanography, 134, 54–76.

Oey, L., Ezer, T. and Lee, H. (2005) Loop Current, rings and related circulation in the Gulf of Mexico: A review of numerical models and future challenges. Geophysical Monograph-American Geophysical Union, 161, 31.

Rooker, J. R., Bremer, J. R. A., Block, B. A., Dewar, H., De Metrio, G., Corriero, A., Kraus, R. T., Prince, E. D., Rodriguez-Marin, E. and Secor, D. H. (2007) Life history and stock structure of Atlantic bluefin tuna (Thunnus thynnus). Reviews in Fisheries Science, 15, 265–310.

Rooker, J. R., Kitchens, L. L., Dance, M. A., Wells, R. D., Falterman, B. and Cornic, M. (2013) Spatial, temporal, and habitat-related variation in abundance of pelagic fishes in the Gulf of Mexico: potential implications of the Deepwater Horizon oil spill. PloS one, 8, e76080.

Rooker, J. R., Simms, J. R., Wells, R. D., Holt, S. A., Holt, G. J., Graves, J. E. and Furey, N. B. (2012) Distribution and habitat associations of billfish and swordfish larvae across mesoscale features in the Gulf of Mexico. PloS one, 7, e34180.

Rost, B., Zondervan, I. and Wolf-Gladrow, D. (2008) Sensitivity of phytoplankton to future changes in ocean carbonate chemistry: current knowledge, contradictions and research directions. Marine Ecology-Progress Series, 373, 227–237.

Saint-Béat, B., Vézina, A. F., Asmus, R., Asmus, H. and Niquil, N. (2013) The mean function provides robustness to linear inverse modelling flow estimation in food webs: A comparison of functions derived from statistics and ecological theories. Ecological Modelling, 258, 53–64.

Scanlan, D. J. and Post, A. F. (2008) Chapter 24 - Aspects of Marine Cyanobacterial Nitrogen Physiology and Connection to the Nitrogen Cycle. In: D. G. Capone, D. A. Bronk, M. R. Mulholland and E. J. Carpenter (eds) Nitrogen in the Marine Environment (2nd Edition). Academic Press, San Diego, pp. 1073–1095.

Schmitz, W., Biggs, D., Lugo-Fernandez, A., Oey, L. Y. and Sturges, W. (2005) A synopsis of the circulation in the Gulf of Mexico and on its continental margins. Circulation in the Gulf of Mexico: Observations and models, 11–29.

Selph, K. E., Landry, M. R., Taylor, A. G., Gutierrez-Rodriguez, A., Stukel, M. R., Wokuluk, J. and Pasulka, A. (2016) Phytoplankton production and taxon-specific growth rates in the Costa Rica Dome. Journal of Plankton Research, 38, 199–215.

Selph, K. E., Swalethorp, R., Stukel, M. R., Knapp, A. N. and Landry, M. R. (in prep.) Phytoplankton assemblages in the open ocean water of the Gulf of Mexico during May 2017 and 2018. Journal of Plankton Research.

Shiroza, A., Malca, E., Gerard, T., Lamkin, J., Landry, M. R., Laiz-Carrión, R., Stukel, M. R. and Swalethorp, R. (in prep.) Diet and prey selection in Atlantic Bluefin Tuna (Thunnus thynnus) larvae within its Gulf of Mexico spawning grounds. Journal of Plankton Research.

Shropshire, T. A., Morey, S. L., Chassignet, E., Coles, V. J., Fiksen, O., Gerard, T., Malca, E., Laiz-Carrión, R., Lamkin, J., Reglero, P., Shiroza, A. and Stukel, M. R. (in prep.) Mortality of Atlantic Bluefin tuna larvae in the Gulf of Mexico due to starvation and predation Journal of Plankton Research.

Shropshire, T. A., Morey, S. L., Chassignet, E. P., Bozec, A., Coles, V. J., Landry, M. R., Swalethorp, R., Zapfe, G. and Stukel, M. R. (2020) Quantifying spatiotemporal variability in zooplankton dynamics in the Gulf of Mexico with a physical-biogeochemical model. Biogeosciences, 17, 3385–3407.

Sigman, D. M., Casciotti, K. L., Andreani, M., Barford, C., Galanter, M. and Bohlke, J. K. (2001) A bacterial method for the nitrogen isotopic analysis of nitrate in seawater and freshwater. Analytical Chemistry, 73, 4145–4153.

Soetaert, K., Van Den Meersche, K. and Van Oevelen, D. (2009) limSolve: Solving linear inverse models R package version. Vol. 1.5.4.

Stoecker, D. K., Hansen, P. J., Caron, D. A. and Mitra, A. (2017) Mixotrophy in the Marine Plankton. Annual Review of Marine Science, 9, 311–335.

Strickland, J. D. and Parsons, T. R. (1972) A practical handbook of seawater analysis, second ed. Bulletin of the Fisheries Research Board of Canada, 167.

Stukel, M. R., Benitez-Nelson, C., Décima, M., Taylor, A. G., Buchwald, C. and Landry, M. R. (2016) The biological pump in the Costa Rica Dome: An open ocean upwelling system with high new production and low export. Journal of Plankton Research, 38, 348–365.

Stukel, M. R., Décima, M. and Kelly, T. B. (2018a) A new approach for incorporating 15N isotopic data into linear inverse ecosystem models with Markov Chain Monte Carlo sampling. PloS one, 13, e0199123.

Stukel, M. R., Décima, M., Landry, M. R. and Selph, K. E. (2018b) Nitrogen and isotope flows through the Costa Rica Dome upwelling ecosystem: The crucial mesozooplankton role in export flux. Global Biogeochemical Cycles, 32, 1815–1832.

Stukel, M. R., Kahru, M., Benitez-Nelson, C. R., Decima, M., Goericke, R., Landry, M. R. and Ohman, M. D. (2015) Using Lagrangian-based process studies to test satellite algorithms of vertical carbon flux in the eastern North Pacific Ocean. Journal of Geophysical Research: Oceans, 120, 7208–7222.

Stukel, M. R., Kelly, T. B., Aluwihare, L. I., Barbeau, K. A., Goericke, R., Krause, J. W., Landry, M. R. and Ohman, M. D. (2019) The Carbon:^234^Thorium ratios of sinking particles in the California Current Ecosystem 1: Relationships with plankton ecosystem dynamics. Marine Chemistry, 212, 1–15.

Stukel, M. R., Landry, M. R., Ohman, M. D., Goericke, R., Samo, T. and Benitez-Nelson, C. R. (2012) Do inverse ecosystem models accurately reconstruct plankton trophic flows? Comparing two solution methods using field data from the California Current. Journal of Marine Systems, 91, 20–33.

Swalethorp, R., Laiz-Carrión, R., Malca, E., Stukel, M. R., Quintanilla, J. M., Gerard, T., Lamkin, J., Shiroza, A. and Landry, M. R. (in prep.) Trophic structure and nitrogen sources supporting Gulf of Mexico Atlantic Bluefin Tuna (Thunnus thynnus) larvae. Journal of Plankton Research.

Taylor, A. G. and Landry, M. R. (2018) Phytoplankton biomass and size structure across trophic gradients in the southern California Current and adjacent ocean ecosystems. Marine Ecology Progress Series, 592, 1–17.

Teo, S. L., Boustany, A., Dewar, H., Stokesbury, M. J., Weng, K. C., Beemer, S., Seitz, A. C., Farwell, C. J., Prince, E. D. and Block, B. A. (2007) Annual migrations, diving behavior, and thermal biology of Atlantic bluefin tuna, Thunnus thynnus, on their Gulf of Mexico breeding grounds. Marine Biology, 151, 1–18.

Tilley, J. D., Butler, C. M., Suárez-Morales, E., Franks, J. S., Hoffmayer, E. R., Gibson, D. P., Comyns, B. H., Ingram Jr, G. W. and Blake, E. M. (2016) Feeding ecology of larval Atlantic bluefin tuna, Thunnus thynnus, from the central Gulf of Mexico. Bulletin of Marine Science, 92, 321–334.

Turner, J. T. (1986) Zooplankton feeding ecology: contents of fecal pellets of the cyclopoid copepods Oncaea venusta, Corycaeus amazonicus, Oithona plumifera, and O. simplex from the northern Gulf of Mexico. Marine Ecology, 7, 289–302.

Uye, S.-I. and Kayano, Y. (1994) Predatory feeding behavior of Tortanus (Copepoda: Calanoida): life-stage differences and the predation impact on small planktonic crustaceans. Journal of Crustacean Biology, 14, 473–483.

Van Den Meersche, K., Soetaert, K. and Van Oevelen, D. (2009) xSample(): An R function for sampling linear inverse problems. Journal of Statistal Software, Code Snippets, 30, 1–15.

Van Oevelen, D., Van Den Meersche, K., Meysman, F. J. R., Soetaert, K., Middelburg, J. J. and Vezina, A. F. (2010) Quantifying food web flows using linear inverse models. Ecosystems, 13, 32–45.

Vézina, A. F. and Platt, T. (1988) Food web dynamics in the ocean .1. Best estimates of flow networks using inverse methods. Marine Ecology Progress Series, 42, 269–287.

Walters, C., Martell, S. J., Christensen, V. and Mahmoudi, B. (2008) An Ecosim model for exploring Gulf of Mexico ecosystem management options: implications of including multistanza life-history models for policy predictions. Bulletin of Marine Science, 83, 251–271.

Wells, R. D., Rooker, J. R., Quigg, A. and Wissel, B. (2017) Influence of mesoscale oceanographic features on pelagic food webs in the Gulf of Mexico. Marine Biology, 164, 92.

Yingling, N., Kelly, T. B., Selph, K. E., Landry, M. R., Knapp, A. N., Kranz, S. A. and Stukel, M. R. (in prep.) Taxon-specific phytoplankton growth, nutrient limitation, and light limitation in the oligotrophic Gulf of Mexico. Journal of Plankton Research.

Zehr, J. P. (2011) Nitrogen fixation by marine cyanobacteria. Trends in Microbiology, 19, 162–173.

